# Distinct subpopulations of ventral pallidal cholinergic projection neurons encode valence of olfactory stimuli

**DOI:** 10.1101/2023.10.06.561261

**Authors:** Ronald Kim, Mala Ananth, Niraj S. Desai, Lorna W. Role, David A. Talmage

## Abstract

The ventral pallidum (VP) mediates motivated behaviors largely via the action of VP GABA and glutamatergic neurons. In addition to these neuronal subtypes, there is a population of cholinergic projection neurons in the VP, whose functional significance remains unclear. To understand the functional role of VP cholinergic neurons, we first examined behavioral responses to an appetitive (APP) odor that elicited approach, and an aversive (AV) odor that led to avoidance. To examine how VP cholinergic neurons were engaged in APP vs. AV responses, we used an immediate early gene marker and in-vivo fiber photometry, examining the activation profile of VP cholinergic neurons in response to each odor. Exposure to each odor led to an increase in the number of cFos counts and increased calcium signaling of VP cholinergic neurons. Activity and cre-dependent viral vectors were designed to label engaged VP cholinergic neurons in two distinct contexts: (1) exposure to the APP odor, (2) followed by subsequent exposure to the AV odor, and vice versa. These studies revealed two distinct, non-overlapping subpopulations of VP cholinergic neurons: one activated in response to the APP odor, and a second distinct population activated in response to the AV odor. These two subpopulations of VP cholinergic neurons are spatially intermingled within the VP, but show differences in electrophysiological properties, neuronal morphology, and projections to the basolateral amygdala. Although VP cholinergic neurons are engaged in behavioral responses to each odor, VP cholinergic signaling is only required for approach behavior. Indeed, inhibition of VP cholinergic neurons not only blocks approach to the APP odor, but reverses the behavior, leading to active avoidance. Our results highlight the functional heterogeneity of cholinergic projection neurons within the VP. These two subpopulations of VP cholinergic neurons differentially encode valence of olfactory stimuli and play unique roles in approach and avoidance behaviors.

## Introduction

Proper decision-making is critical for survival in a dynamic environment with both appetitive (APP) and aversive (AV) stimuli. This entails motivation to approach and retrieve rewarding stimuli, as well as motivation to avoid harmful stimuli. The first step in any behavior is the ability to properly encode valence (Tye, 2018). The valence of the stimulus (either positive or negative) then directs the animal towards an appropriate behavioral output. Generally, animals demonstrate approach behavior towards positive valence stimuli, whereas negative valence stimuli elicit avoidance behaviors (Tye, 2018; Smith and Torregrossa, 2021; Warlow and Berridge, 2021). Misattribution of valence, however, can lead to maladaptive behaviors. Furthermore, prolonged improper valence encoding can lead to the development of psychiatric diseases such as drug addiction and depression (Kalivas and Volkow, 2005; Fox and Lobo, 2019). Accordingly, examining the brain regions and neural circuits that underlie proper valence encoding is essential to understanding mechanisms underlying motivated behaviors.

The ventral pallidum (VP) is involved in mediating motivated behaviors (Root et al., 2015; Stephenson-Jones et al., 2020; Farrell et al., 2021; Lederman et al., 2021). The VP coordinates limbic inputs and regulates motivated behaviors in response to these inputs (Smith et al., 2009; Root et al., 2015). The VP has been implicated in a variety of neuropsychiatric disorders that are characterized by motivational imbalance, including drug addiction, depression, stress, and anxiety (Gardner, 2011; Knowland et al., 2017; McGovern and Root, 2019; Liu et al., 2020; Ji et al., 2023).

The VP modulates motivation via the activity of multiple different cell-types. GABAergic projection neurons in the VP increase motivation to receive a rewarding stimulus (Faget et al., 2018; Heinsbroek et al., 2020; Stephenson-Jones et al., 2020). For example, optogenetic stimulation of VP GABA neurons makes a stimulus appear more rewarding and can drive reinforcement behavior (Faget et al., 2018; Stephenson-Jones et al., 2020). In contrast, stimulation of glutamatergic projection neurons in the VP enhances the aversive responses, leading to avoidance (Faget et al., 2018; Stephenson-Jones et al., 2020).

In addition to GABA and glutamate neurons, the VP also includes a small population of cholinergic neurons that project out of the VP to targets including the basolateral amygdala (BLA), medial prefrontal cortex and thalamus (Root et al., 2015; Faget et al., 2018; Záborszky et al., 2018). The functional significance of the VP cholinergic projection neurons is unclear. Using electrophysiology recordings, Stephenson-Jones and colleagues identified a cluster of VP neurons, described as Type 1 neurons, whose firing patterns differed from that of either VP GABAergic or glutamatergic neurons (Stephenson-Jones et al., 2020). Unlike VP GABA or glutamatergic neurons, these neurons responded to both aversive and rewarding stimuli (Stephenson-Jones et al., 2020). These Type I VP neurons were hypothesized to be salience encoding cholinergic neurons since their activity patterns resemble those of cholinergic neurons in other parts of the basal forebrain, which respond to both reward and punishment (Hangya et al., 2015).

Despite the known presence of cholinergic projection neurons in the VP, the functional significance of these neurons remains unknown. The previously identified Type I neurons of the VP (Stephenson-Jones et al., 2020), indicate VP cholinergic neurons could respond to both positive and negative valence stimuli. However, it is unknown if the same neuron responds to each stimulus (thus encoding salience), or if distinct subsets of VP cholinergic neurons are uniquely activated in response to positive vs. negative valence stimuli (therefore encoding valence). Accordingly, we utilized activity- and cre-dependent viral vectors to permanently label activated VP cholinergic neurons in distinct behavioral contexts. We examined the activation profile of VP cholinergic neurons following exposure to an APP odor (2-phenylethanol) and compared this profile to that stimulated by an AV odor (predator odor). Our results reveal diverse functional profiles of APP vs. AV cholinergic neurons in the VP. Here, we demonstrate that the VP includes two distinct and non-overlapping subpopulations of cholinergic neurons: one activated in response to an APP odor, and a second, distinct subpopulation activated in response to an AV odor. These two subpopulations of VP cholinergic projection neurons are spatially intermingled within the VP but are differentiated from one another in a variety of characteristics including electrophysiological properties, overall neuronal morphology, and projections to downstream brain regions. Importantly, although VP cholinergic neurons are engaged in APP and AV behavioral responses, VP cholinergic signaling is only required for approach behavior.

## Results

### APP and AV odors elicit innate behavioral responses

To examine if VP cholinergic neurons were engaged in APP and/or AV behaviors, we first measured behavioral responses to an APP odor (2-phenylethanol) and to an AV odor (predator odor, mountain lion urine). We directly quantified preference for each odor in a two-arm preference test (Y-Maze: Fig 1A). In a preference test between saline (null odor, N) and the APP odor, mice spent significantly more time in the arm with the APP odor (*t* (16) = −5.7, *p* < 0.001; Fig 1B and 1C), indicating approach to the APP odor. In contrast, in a preference test between saline and the AV odor, mice spent significantly more time in the arm with saline (*t* (14) = −5.06, *p* < 0.001; Fig 1B and 1C), indicating avoidance of the AV odor. Neither the APP nor the AV odor affected total distance traveled or velocity (Fig S1).

**Figure 1:**
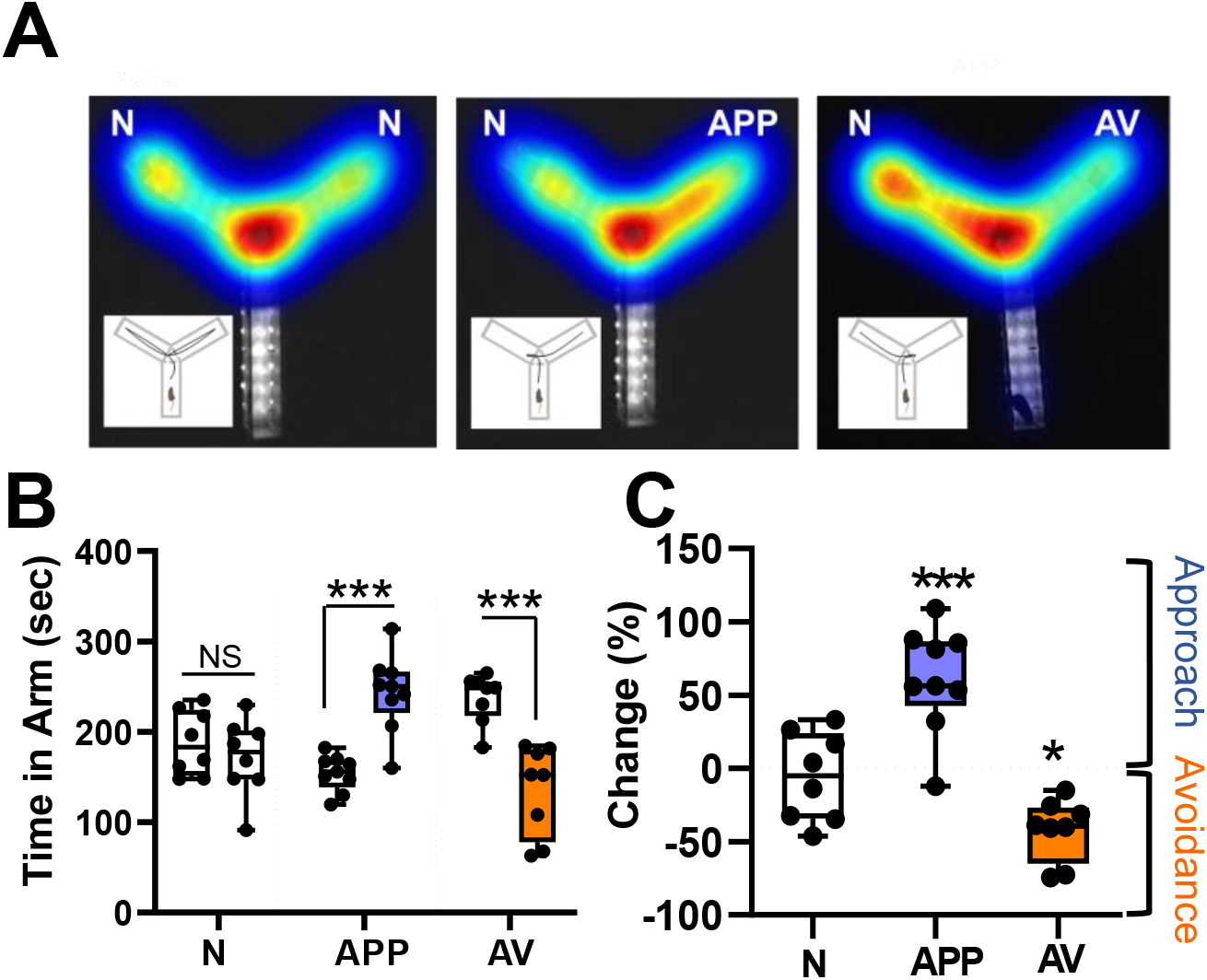
Appetitive odors elicit approach, whereas aversive odors elicit avoidance behaviors. A. Example heatmaps from each of the behavioral paradigms tested *(from left to right):* Null Odor (N, saline diluent), Appetitive Odor (APP, 2-phenylethanol), Aversive Odor (AV, mountain lion urine). Insets illustrate the path traveled by typical mouse under each condition (N, APP, AV). Left: Presentation of saline in both test arms (N) results in approximately equal time spent in each arm. Middle: Presentation of an appetitive odor (APP) in one arm elicits approach behavior, defined as more time spent in the arm with the APP odor than in the arm with null odor (N). Right: Presentation of an aversive odor (AV) elicits avoidance behavior, defined as more time spent in the arm with null odor (N) than in the AV arm. B. Left: Total time spent in each arm under N vs N conditions (8 mice), compared with N vs APP (9 mice), and N vs AV (8 mice) conditions. Mice spent significantly more time in the APP vs N arm (approach) and spent significantly less time in the AV vs N arm (avoidance). *** *p* <0.001. C. Behavioral responses measured as percent change of time spent in test vs N arm. Mice spent significantly more time in the arm with the APP odor and significantly less time with the AV odor compared to time spent in N arm. All results were independent of whether left or right arm was used as test arm. *** *p* <0.001; * *p*< 0.05.

### Approach and avoidance behaviors engage VP cholinergic neurons

To test if immediate early gene expression (cFos) was increased in VP cholinergic neurons following odor exposure, mice were euthanized 45-50 min following the odor preference test (Fig 1), and fixed tissue slices containing the VP were stained with antibodies recognizing ChAT (cholinergic neurons) and cFos (as a marker of neuronal activation) (Fig 2A and 2B). The number of cholinergic neurons identified was equivalent between groups (ChAT counts, Fig 2C left). The total number of cFos+ cells in the VP significantly increased following exposure to either odor (*F* (2, 64) = 12.79, *p* < 0.001; pairwise comparisons: APP vs. saline (*t* = 4.29, *p* < 0.001), AV vs. saline (*t* = 4.64, *p* < 0.001); Fig 2C right). Importantly, cholinergic neurons were activated following exposure to either odor (number of colocalized ChAT and cFos+ neurons (Fig 2D left: *F* (2, 64) = 4.19, *p* < 0.05; pairwise comparison: APP vs. saline (*t* = 2.77, *p* < 0.05), AV vs. saline (*t* = 2.26, *p* < 0.05)); percentage of ChAT neurons that were cFos+ after odor exposure (Fig 2D right: (*F* (2, 64) = 4.35, *p* < 0.05; pairwise comparisons: APP vs. saline (*t* = 2.63, *p* < 0.05), AV vs. saline (*t* = 2.59, *p* < 0.05)).

**Figure 2:**
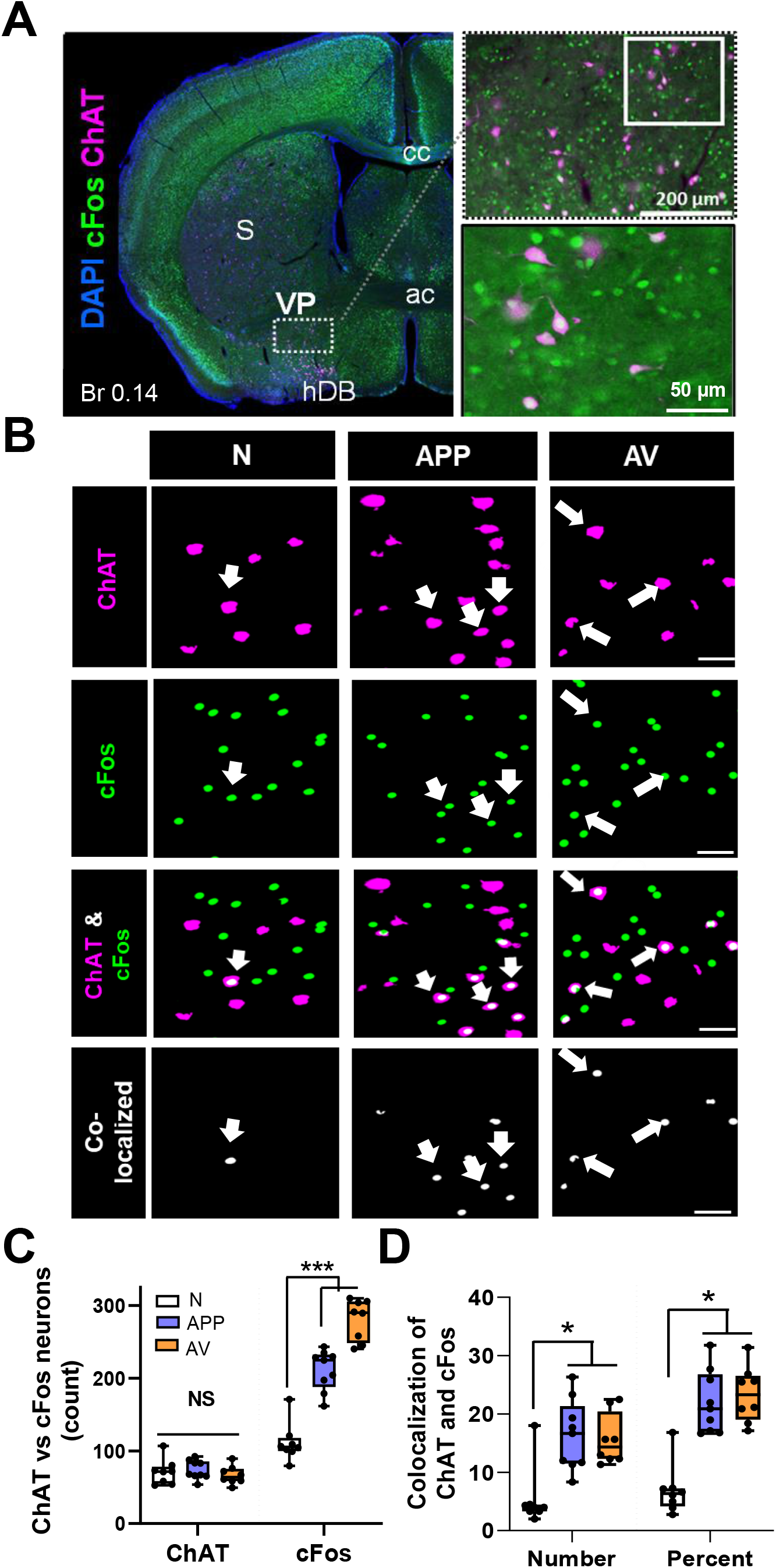
Appetitive and aversive odors increase activation of cholinergic neurons within the VP as indicated by an increase of the immediate early gene cFos. **A.** Left: Representative image of activated cholinergic neurons in a coronal slice (Br 0.14) following odor preference test. The VP is defined as the sparse cluster of cholinergic neurons ventral to the striatum (S), ventrolateral to the anterior commissure (ac), and dorsolateral to the dense cluster of cholinergic neurons that comprise the horizontal limb of the diagonal band (hDB). Right: Higher magnification images of the VP region (top), with additional magnification of the demarcated area in the inset below. Blue = DAPI, Green = cFos, Magenta = ChAT. **B.** Representative high magnification images of the VP following ChAT and cFos immunohistochemistry from mice that underwent an odor preference test ∼45 min prior to euthanasia and tissue processing. The first column shown are “null” odor presentation (n = 8). The 2^nd^ column shown are from mice following an appetitive odor preference test (APP, n = 9). The 3^rd^ column shown are from mice post an aversive odor preference test (AV, n = 8). Rows are representative images for ChAT (1^st^ row), cFos (2^nd^ row), the overlay between ChAT and cFos (3^rd^ row) and colocalization between ChAT and cFos (bottom row). Scale bar is 50 µm. Arrowheads represent colocalized ChAT and cFos neurons in the VP. **C.** Total counts of neurons immunostained for both ChAT and cFos. Mice exposed to either odor (APP or AV) showed a significant increase in total cFos+ cells in the VP (right).*** *p* < .001. Note: no significant differences between groups in the total number of ChAT neurons assayed (left). **D.** Colocalization of ChAT and cFos in the VP in total numbers (left) and percentage of ChAT+ neurons that co-express cFos (right). Both odors (APP or AV) significantly increased the number and percentage of VP neurons that expression both ChAT and cFos .*** *p* < .001, * *p* < 0.05.

### VP cholinergic neurons display time-locked increases in calcium activity in direct response to each odor

To examine the activity of VP cholinergic neurons in direct response to each odor, we used Cre-dependent GCaMP6F and in-vivo fiber photometry to measure calcium responses of VP cholinergic neurons during timed delivery of each odor. Mice were exposed to either the APP or AV odor 3 times for 10 seconds, with a 3-minute interval between exposures, while we continuously recorded GCaMP signals (Fig 3A). Approximately 24-hours later, the same mice were exposed to the opposite valence odor (counter-balanced to odor exposure). Both the APP odor (Fig 3B, 3D and 3F) and the AV odor (Fig 3C, 3E and 3G) increased calcium activity of VP cholinergic neurons during the 10-second delivery of each odor. The area under the curve (AUC) was significantly increased during each APP odor delivery (*F* (4, 34) = 14.28, *p* < 0.001; Fig 3D, pairwise comparisons: APP 1 vs. pre-odor (*t* (6.35, *p* < 0.001), APP 1 vs. post-odor (*t* (5.66, *p* < 0.001), APP 2 vs. pre-odor (*t* (3.02, *p* < 0.05), APP 3 vs. pre-odor (*t* (4.74, *p* < 0.001), APP 3 vs. post-odor (*t* (4.04, *p* < 0.05)), as well as following each AV odor delivery (*F* (4, 34) = 12.34, *p* < 0.001; Fig 3E, pairwise comparisons: AV 1 vs. pre-odor (*t* (5.78, *p* < 0.001), AV 1 vs. post-odor (*t* (5.15, *p* < 0.001), AV 2 vs. pre-odor (*t* (4.16, *p* < 0.05), AV 2 vs. post-odor (*t* (3.53, *p* < 0.05), AV 3 vs. pre-odor (*t* (3.81, *p* < 0.05), AV 3 vs. post-odor (*t* (3.19, *p* < 0.05)). The max z-score ΔF/F was significantly increased during the first APP odor (*F* (4, 34) = 4.4, *p* < 0.05, *t* = 3.69, *p* < 0.05; Fig 3F) and first AV odor (*F* (4, 34) = 5.19, *p* < 0.05, *t* = 3.90, *p* < 0.05; Fig 3G). These results corroborate our IHC results and confirm that both APP and AV odors directly engage VP cholinergic neurons.

**Figure 3:**
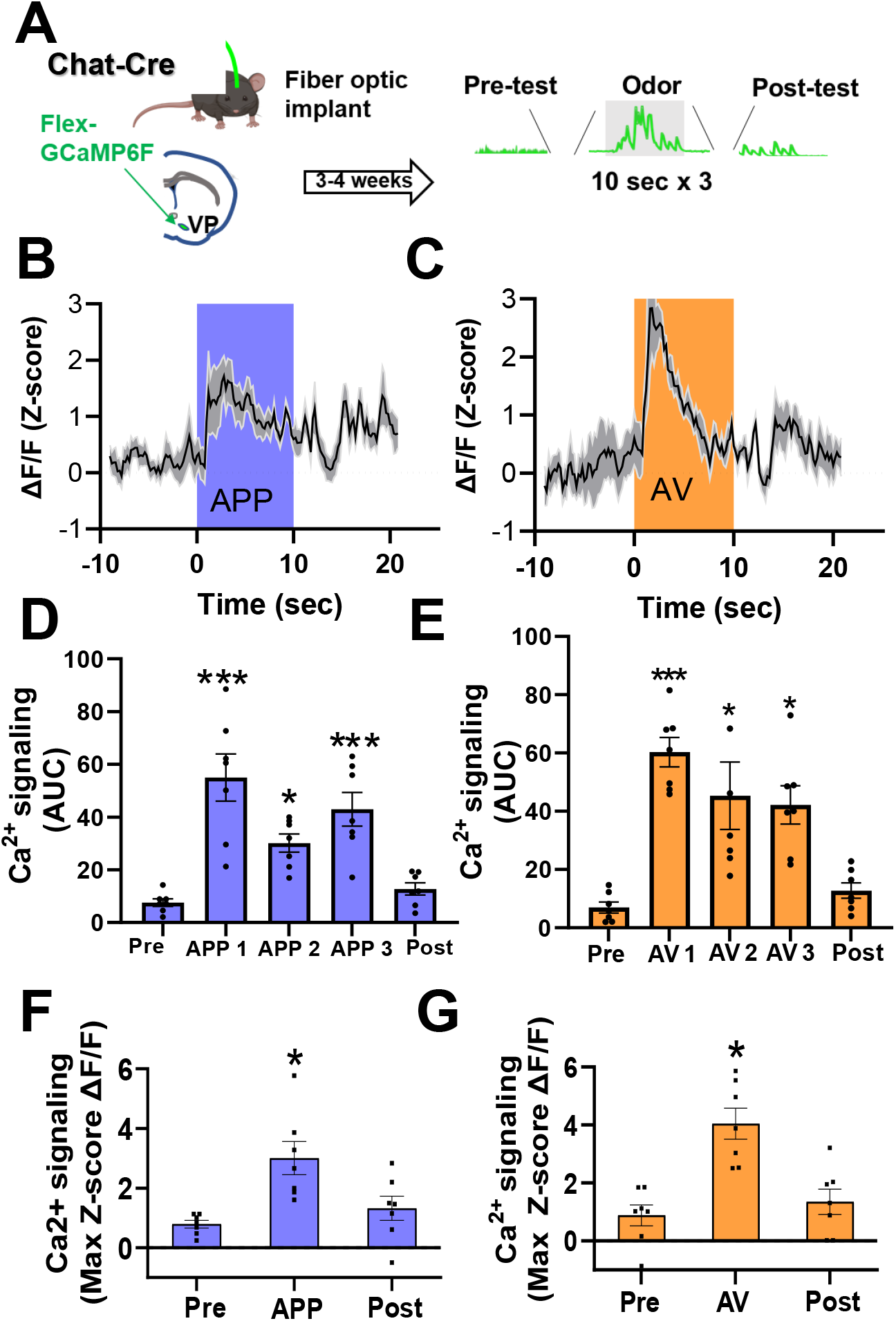
VP cholinergic neurons display time-locked increases in calcium activity in direct response to the appetitive odor and aversive odor. **A.** Workflow and timeline for in-vivo fiber photometric assays of calcium signaling in VP cholinergic neurons during odor presentation. Chat-Cre mice were injected with AAV.Syn.FLEX.GCaMP6F in the VP. Following recovery from surgery (3 - 4 weeks), mice were exposed to timed delivery of either the appetitive (APP) or aversive odor (AV) on Day 1 (3 discreet odor puffs lasting 10 seconds, 3 minutes in between each odor delivery). Approximately 24-hours later, each mouse was exposed to the opposite odor (counter-balanced to odor exposure, n = 7). **B.** VP FLEX-GCaMP6F traces in response to the first 10-second delivery of the APP odor (n = 7). The blue shaded area represents the time window for APP odor delivery. **C.** VP FLEX-GCaMP6F traces in response to the first 10-second delivery of the AV odor (n = 7). The orange shaded area represents the time window for AV odor delivery. **D.** Area under the curve (AUC) measurements before, during and following each APP odor delivery. Compared to pre-test values, the AUC was significantly greater following each APP odor delivery. *** *p* < 0.001, * *p* < 0.05. **E.** AUC measurements before, during and following AV odor exposure. Compared to pre-test and post-test values, the AUC was significantly greater following each AV odor exposure. *** *p* < 0.001, * *p* < 0.05. **F.** The max z-score score ΔF/F before, during and following APP odor exposure (* *p* < 0.05). The max z-score ΔF/F was significantly increased following the first APP odor delivery. * *p* < 0.05. **G.** The max z-score ΔF/F before, during and following AV odor exposure. The max z-score ΔF/F was significantly increased following the first AV odor delivery. * *p* < 0.05.

### Silencing VP cholinergic neurons not only abolishes, but reverses approach behavior to active avoidance

To test the requisite participation of VP cholinergic neurons in approach and/or avoidance behaviors, we used a Cre-dependent inhibitory DREADD to silence cholinergic neurons in the VP. Mice were injected with AAV.Syn.eGFP (sham), or AAV.Syn.eGFP and AAV.hSyn.DIO.hM4Di concurrently (Fig 4A). Following recovery from surgery, mice were injected IP with 0.1 mg/kg clozapine and 15-minutes later underwent behavioral testing, identical to the 10-minute odor preference test described above in Fig 1. In a preference test with the APP odor, mice in the sham group (eGFP only + clozapine) exhibited approach to the APP odor (Fig 4B left and 4C). hM4di inhibition of VP cholinergic neurons abolished this effect and mice spent significantly more time in the saline arm vs. the arm with the APP odor (*t* (8) = 4.21, *p* < 0.05; Fig 4B right and 4C), indicating active avoidance of the APP odor. This indicates that VP cholinergic signaling is required for approach behavior, and inhibition of cholinergic neurons in the VP leads to avoidance of an APP stimulus.

**Figure 4:**
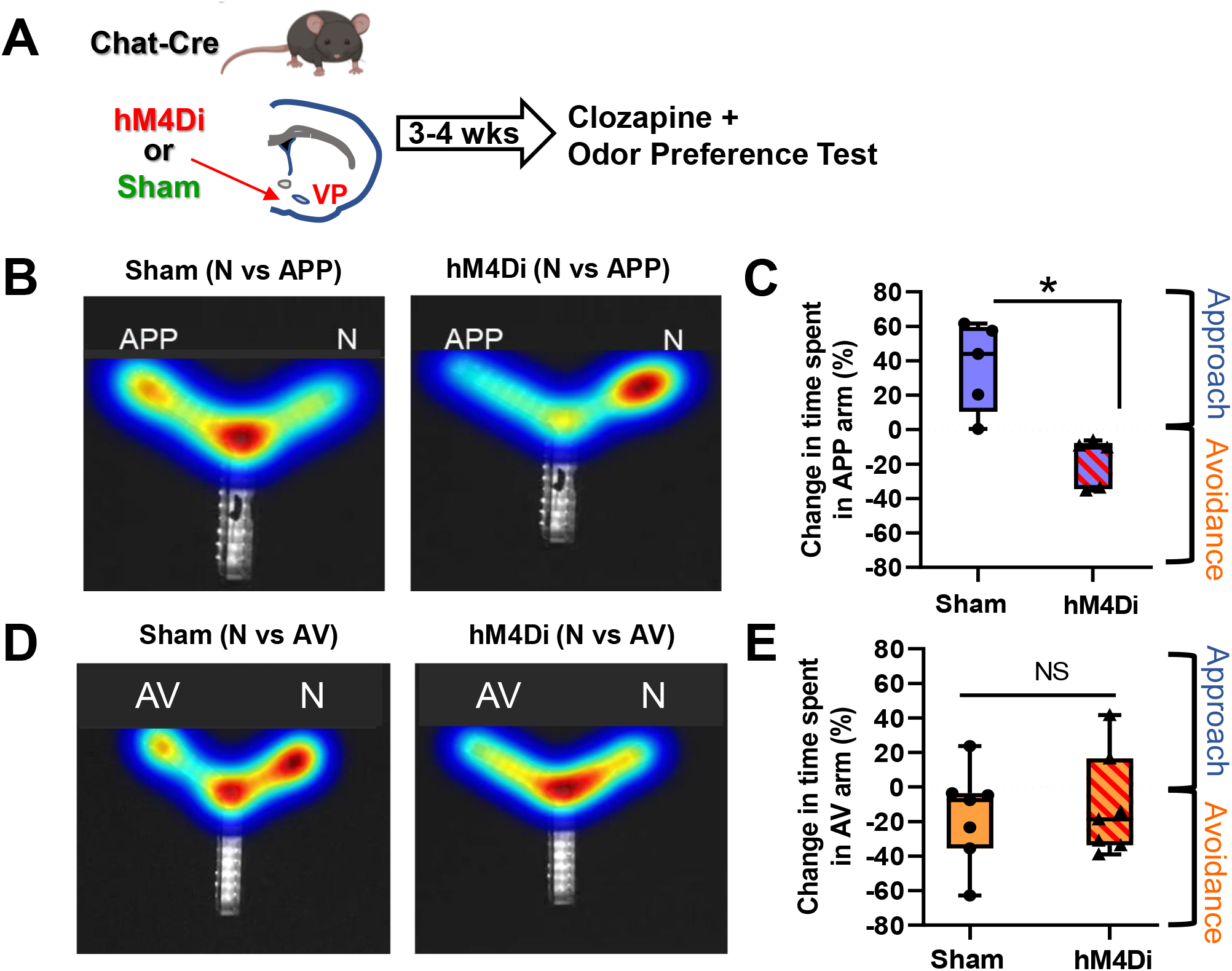
Chemogenetic inhibition of VP cholinergic neurons abolishes approach behavior. **A.** Workflow and timeline of experiments using chemogenetic inhibition to test the contribution of VP cholinergic activity on approach and avoidance behavior. Chat-Cre mice were injected with AAV.Syn.eGFP (Sham), or with AAV.Syn.eGFP and AAV.hSyn.DIO.hM4Di (hM4Di) in the VP. Following recovery from surgery (∼3 - 4 weeks) mice were injected with 0.1 mg/kg clozapine 15 minutes before an odor preference test. Preference for the appetitive odor (APP) or aversive odor (AV) was assessed in a Y-Maze. **B.** Heatmap illustrating the effects of chemogenetic inhibition of VP cholinergic neurons on approach to the APP odor. Left: representative heatmap depicting approach to the APP odor in a sham mouse (eGFP only + 0.1 mg/kg clozapine). Right: Representative heatmap in a mouse with chemogenetic inhibition of the VP (hM4Di) and APP odor presentation. Not only does chemogenetic inhibition of VP cholinergic neurons block approach behavior, but it also reverses the response to a clear avoidance behavior. **C.** Quantification of experiments shown in **(B)** Mice in the sham group (n = 5) exhibit approach to the APP odor. Mice with hM4Di inhibition of VP cholinergic neurons (n = 5) spend more time in the saline paired arm (vs. APP), indicating avoidance of the APP odor. * *p* < 0.05. **D.** Heatmap illustrating the effects of chemogenetic inhibition of VP cholinergic neurons on avoidance of the AV odor. Left: Representative heatmap depicting avoidance of the AV odor in a sham mouse. Right: Representative heatmap showing that inhibition of VP cholinergic neurons is without effect on avoidance behavior. **E.** Quantification of experiments shown in **(D)** Mice in the sham group (n = 7) exhibit innate avoidance of the AV odor. However, inhibition of VP cholinergic neurons had no effect on avoidance of the AV odor. Mice with hM4Di inhibition of VP cholinergic neurons (n = 7) exhibit a similar degree of avoidance of the AV odor as sham operated mice.

In contrast to the dramatic effects of hM4Di inhibition of VP cholinergic neurons on the reversal of approach behavior, the behavioral responses towards the AV odor was resistant to VP cholinergic inhibition. In a preference test with the AV odor, mice in the sham group exhibited avoidance of the AV odor (Fig 4D left and 4E). Mice with hM4Di inhibition of VP cholinergic neurons also displayed avoidance of the AV odor (Fig 4D right and 4E). These results indicate inhibition of cholinergic signaling in the VP is insufficient to alter avoidance behavior. These results were not due to any hM4di induced changes in behavior as mice in both groups displayed comparable locomotor activity (Fig S3).

### Distinct and non-overlapping subpopulations of VP cholinergic neurons are activated in response to APP vs. AV odors

Exposure to either odor led to equivalent numbers (∼20%) of VP cholinergic neurons colocalized with cFos (Fig 2) and resulted in a similar increase in calcium activity (Fig 3). To determine if the same or distinct subpopulations of VP cholinergic neurons were activated in response to each odor, we injected the VP of Chat-Cre x Fos-tTA/GFP mice, with AAV_9_.DIO.TRE.hM4Di.P2A.mCherry (ADCD, (Rajebhosale et al., 2021)). In activated cholinergic neurons, tTA will drive hM4Di and mCherry expression when mice are off doxycycline containing food (DOX off). Injected mice were switched from a DOX on to DOX off diet, exposed to either saline, APP or AV odor and then put back on DOX chow. Twenty-four hours later mice were exposed to the same or different odor and then processed for ChAT (to mark cholinergic neurons) and GFP (to amplify the cFos-GFP signal) IHC (Fig 5A).

**Figure 5:**
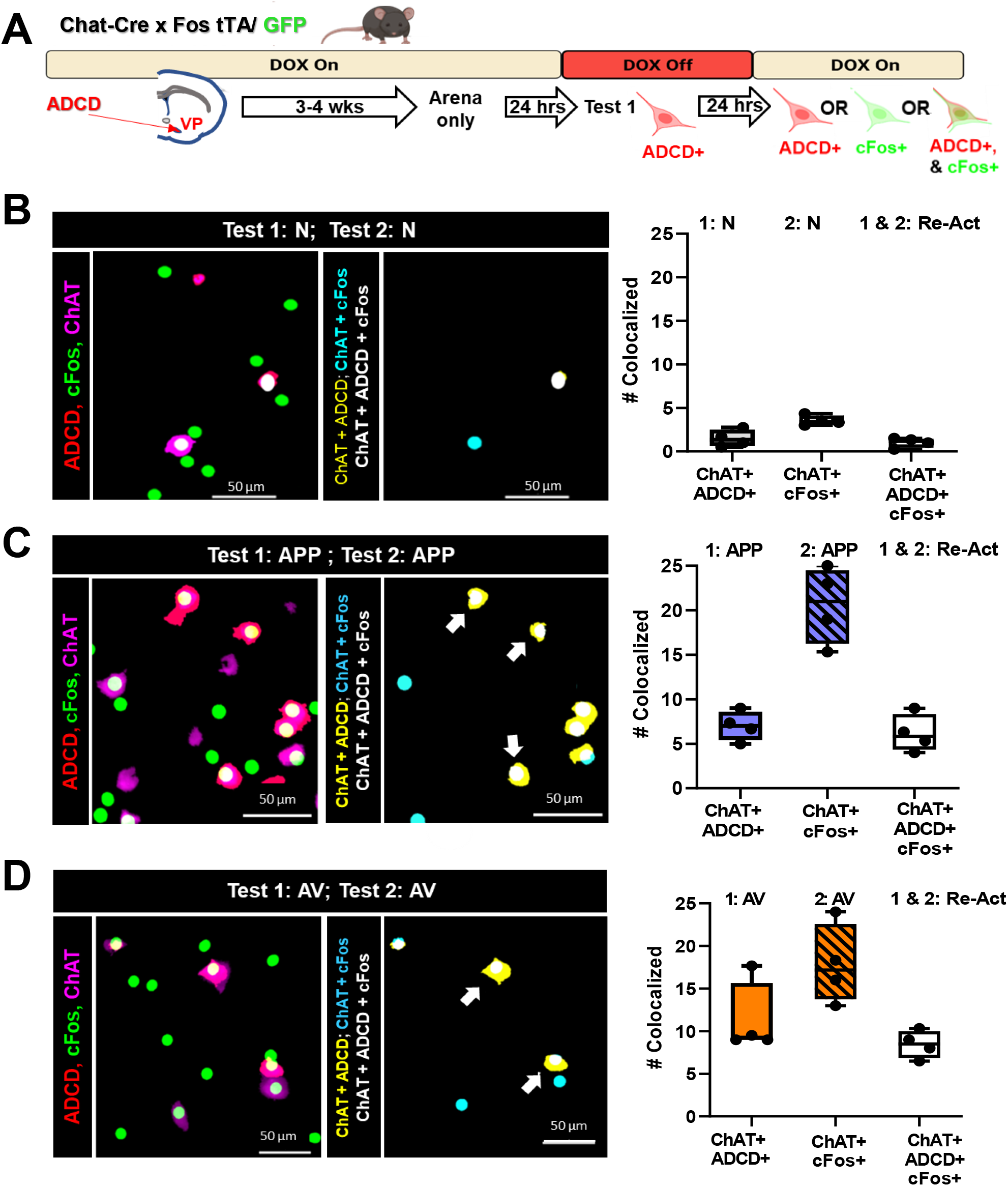
Assessment of activated and reactivated VP cholinergic neurons using genetic and IEG probes following repeat odor presentation. **A.** Schematic diagram of the strategy employed to differentially label VP cholinergic neurons that were activated in distinct contexts. Chat-Cre x cFos tTA/GFP mice were injected with an activity- and cre- dependent (AAV_9_.DIO.TRE.hM4Di.P2A.mCherry (ADCD), Rajebhosale, Ananth et al, 2021) construct in the VP. Following recovery (3 - 4 weeks), mice underwent a behavioral paradigm that lasted 3 days. On day 0, mice were habituated (2 x 10 min) within the Y-Maze arena. Following arena habituation, DOX diet was removed and replaced with regular chow to allow ADCD expression. After 24 hours, mice were exposed to Test 1 conditions in one arm of the Y-Maze with the other arm blocked, thus labeling VP cholinergic neurons activated during Test 1 conditions with ADCD. Mice were then returned to a DOX diet to block further ADCD expression. After a subsequent 24-hour interval, mice were exposed to Test 2 conditions in a different arm of the Y-Maze (with complimentary arm blocked). Mice were euthanized 2.5 hours later for tissue processing for ChAT and cFos IHC. **B.** Mapping (left) and quantification (right) of the activation and reactivation profile of VP cholinergic neurons following 2 exposures to an odor-null stimulus (N, saline diluent) with a 24-hour inter-exposure interval. **Left:** Representative image of ADCD activated VP cholinergic neurons (red), cFos activated neurons (green) and ChAT (cholinergic marker; magenta) following 2 tests with the same null odor stimulus. Mapping of the co-localization of the indicated probes is shown in the right image. **Right:** Quantification of the number of VP cholinergic neurons activated by Test 1 (ChAT+ and ADCD+), those activated by Test 2 (ChAT+ and cFos+) and the neurons that were reactivated (ChAT+, ADCD + and cFos +). Neutral conditions elicits the activation (and reactivation) of very few VP cholinergic neurons. **C.** Mapping (left) and quantification (right) of the activation and reactivation profile of VP cholinergic neurons following 2 test exposures to the same appetitive odor (APP) with a 24-hour inter-test interval. **Left:** Representative images depicting ADCD activated VP cholinergic neurons (red), cFos activated neurons (green) and ChAT (magenta) following 2 tests with the APP odor. Mapping of the co-localization of the indicated probes is shown in the right image right. **Right:** Quantification of the number of VP cholinergic neurons activated by Test 1 (ChAT+ and ADCD+) those activated by Test 2 (ChAT+ and cFos+) and the neurons that were reactivated (ChAT+, ADCD + and cFos +). The APP odor elicits the activation of 5-10 x more VP cholinergic neurons than activated by the neutral stimulus, with Test 2 eliciting an even larger response. Note that all the neurons activated by the first exposure to the APP odor were reactivated by the second exposure to the same APP odor 24 hours later (Re-Act). **D.** Mapping (left) and quantification (right) of the activation and reactivation profile of VP cholinergic neurons following 2 test exposures to the same aversive odor (AV) with a 24-hour inter-test interval. **Left:** Representative images depicting ADCD activated VP cholinergic neurons (red), cFos activated neurons (green) and ChAT (magenta) following 2 tests with the aversive odor. Mapping of the co-localization of the indicated probes is shown in the right image. **Right:** Quantification of the number of VP cholinergic neurons activated by Test 1 (ChAT+ and ADCD+), those activated by Test 2 (ChAT+ and cFos+) and the neurons that were reactivated (ChAT+, ADCD + and cFos +). The AV odor elicits the activation of 10-20 x more VP cholinergic neurons than activated by the neutral stimulus, again with Test 2 eliciting an even larger response than Test 1. Nearly all the neurons activated by the first exposure to the aversive odor were reactivated by the second exposure to the same AV odor 24 hours later (Re-Act).

Very few ADCD+ or cFos+ cholinergic VP neurons were detected in animals exposed and re-exposed to saline (N, Fig 5B). Both the first exposure and the re-exposure to either the APP or the AV odor activated VP cholinergic neurons (ChAT+/ADCD+, ChAT+/cFos+; Fig 5C & 5D). Notably, in each case, virtually all of the ADCD+ VP cholinergic neurons (activated during the first exposure) were also cFos+ (re-activated during the second exposure).

In a distinct cohort of animals, we switched the second odor presented so that it was distinct from the first odor (i.e., from APP to AV, or vice versa: Fig 6). In these mice, we found that similar numbers of VP cholinergic neurons were activated during the first odor exposure (ChAT+/ADCD+, Fig 6B & 6C) and during the second odor exposure (ChAT+/cFos+, Fig 6B & 6C) as seen in the previous cohort (compare Fig 5C and 5D with Fig 6B and 6C). However, in these mice, there were essentially no reactivated VP cholinergics neurons regardless of which odor was presented first (i.e., no ADCD+/cFos+ VP cholinergic neurons).

**Figure 6:**
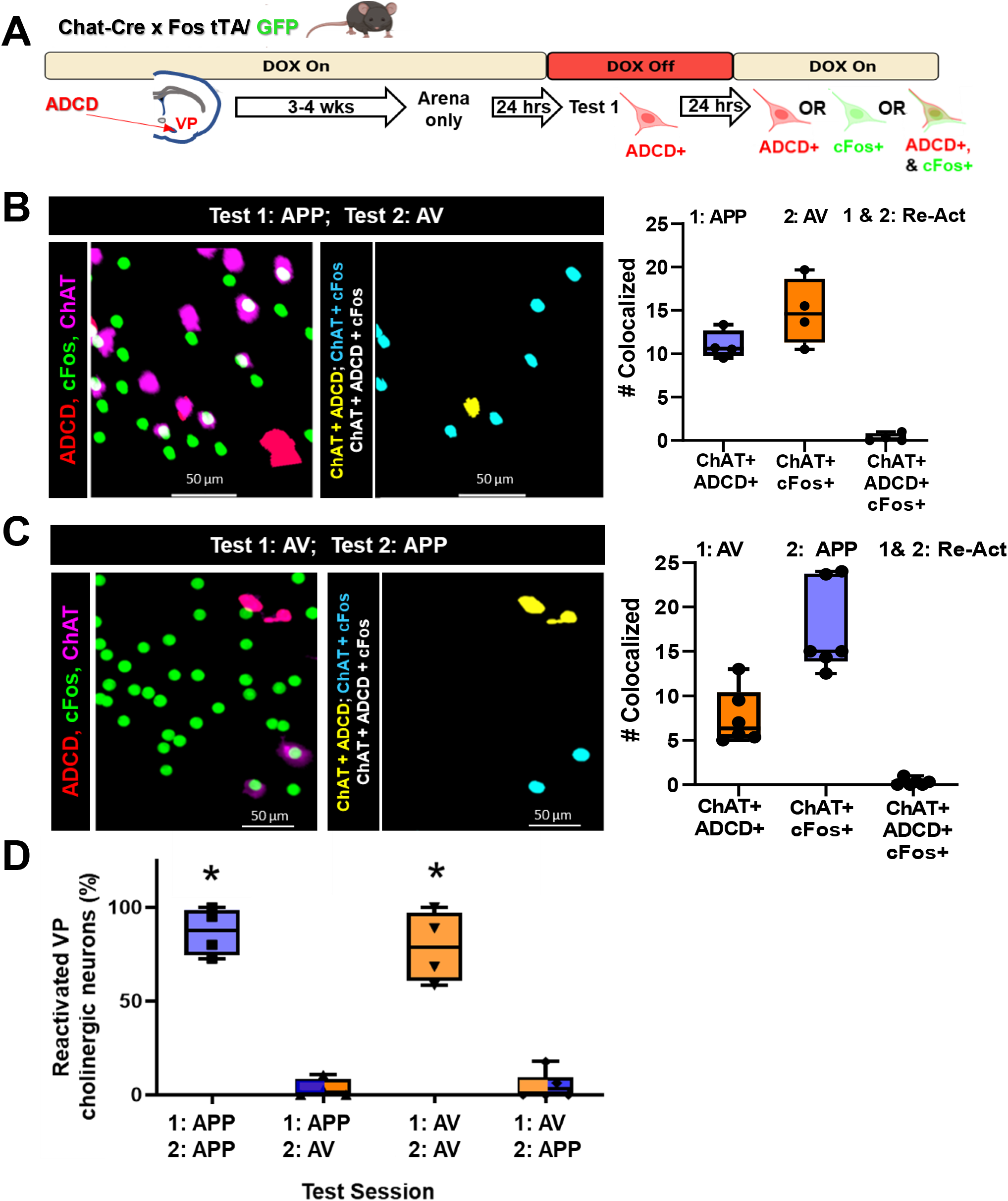
Distinct populations of VP cholinergic neurons are activated in response to distinct olfactory stimuli. **A.** Schematic diagram of the strategy employed to label activated VP cholinergic neurons in distinct contexts (see Figure 4 legend for details). **B.** Mapping (left) and quantification (right) of the activation and reactivation profile of VP cholinergic neurons following an initial exposure to the appetitive odor (Test 1: APP), followed 24-hours later with exposure to the aversive odor (Test 2: AV). **Left:** Representative images showing ADCD activated VP cholinergic neurons (red), cFos activated neurons (green) and ChAT (cholinergic marker; magenta) following an initial test with the APP odor and a second exposure 24 later to the AV odor. Note the lack of colocalized (ChAT+, ADCD+ and cFos+), reactivated neurons in the image map shown on the right. **Right:** Quantification of the number of VP cholinergic neurons activated by Test 1 (ChAT+ and ADCD+) those activated by Test 2 (ChAT+ and cFos+) (Test 1: APP, Test 2: AV). Note that none of the neurons initially activated by the APP odor were also activated by the AV odor (ChAT+, ADCD+ and cFos+). **C.** Mapping (left) and quantification (right) of the activation and reactivation profile of VP cholinergic neurons following an initial exposure to the aversive odor (Test 1: AV), followed by exposure to the appetitive odor (Test 2: APP) with a 24-hour inter-exposure interval. **Left:** Representative images showing ADCD activated VP cholinergic neurons (red), cFos activated neurons (green) and ChAT (cholinergic marker; magenta) following an initial test with AV and a second exposure 24 hours later to APP. Note the complete lack of colocalized (ChAT+, ADCD+ and cFos+), reactivated neurons in the image map shown on the right. **Right:** Quantification of the number of VP cholinergic neurons activated by Test 1 (ChAT+ and ADCD+) those activated by Test 2 (ChAT+ and cFos+) (Test 1: APP, Test 2: AV). Note that none of the neurons initially activated by the AV odor were also activated by the APP odor (ChAT+, ADCD+ and cFos+). **D.** Summary of the quantification of the reactivation profile of VP cholinergic neurons following 2 test exposures to the same odor (either APP or AV) and following exposure to opposite valence odors on Test 1 vs. Test 2. When mice are exposed to the opposite valence odors, there is a complete lack of reactivation, regardless of the order of odor presentation. These data are consistent with APP encoding VP cholinergic neurons being a distinct subpopulation from VP cholinergic neurons encoding AV. * *p* < 0.05.

In sum, on Day 1, each odor significantly increased the number of activated VP cholinergic neurons vs. saline (Fig S4). Likewise, on Day 2, each odor significantly increased the number of activated VP cholinergic neurons vs. saline (Fig S4). However, the number of re-activated VP cholinergic neurons (ChAT+/ADCD+/cFos+ triple positive neurons) was dependent on which odor was presented on Day 2. If mice were exposed to the same odor on Days 1 and 2, there was a significantly greater number of re-activated VP cholinergic neurons (*F* (3, 17) = 68.8, *p* < 0.001; Fig 6D, APP/APP and AV/AV compared to AP/AV and AV/AP), whereas if mice were exposed to a different odor on Day 2, there was no overlap between ChAT+/ADCD+ and ChAT+ /cFos+ VP cholinergic neurons. This indicates VP cholinergic neurons that were activated in response to the first odor are not re-activated when mice are exposed to a distinct odor on Day 2. We replicated these findings using the robust activity marking (RAM) system in C57 mice (Fig S5 and S6). These findings indicate that there are distinct and non-overlapping subpopulations of VP cholinergic neurons that are activated and reactivated in response to APP or AV odors.

### Selective inhibition of VP cholinergic neurons previously activated by the APP odor subpopulations of VP cholinergic neurons abolishes and reverses approach behavior

Our experiment using an inhibitory DREADD for general inhibition of VP cholinergic neurons showed that although VP cholinergic neurons are engaged in response to both the APP and AV, VP cholinergic signaling is only required for approach behavior. To determine the participation of each specific subpopulation of VP cholinergic neurons in approach and/or avoidance behaviors, we used ADCD to selectively silence VP cholinergic neurons that were previously activated by each odor (Fig 7A). Chat-cre x Fos-tTA/GFP mice were injected with ADCD and/or AAV.Syn.eGFP in the VP and underwent behavioral testing identical to the methods described above in the ADCD labeling experiments (Fig 7A). Following recovery from surgery, mice were habituated in the Y-Maze and removed from a DOX diet. Approximately 24-hours later, mice were exposed to an odor (either APP or AV) in one arm of the Y-Maze and returned to a DOX diet. VP cholinergic neurons activated in response to the odor were thus selectively labeled with ADCD. The next day, mice were injected with 0.1 mg/kg clozapine and odor preference was measured in two-arm choice preference test. As expected, in a preference test with the APP odor, mice in the sham group (eGFP only + clozapine) displayed the usual approach behavior in response to the APP odor (Fig 7B left and 7C). However, selective silencing of VP cholinergic neurons that were previously activated by exposure to the APP odor abolished this behavior (*t* (11) = 2.72, *p* < 0.05; Fig 7B right and 7C). Particularly striking was that inhibition of previously activated APP odor VP cholinergic neurons not only blocked approach behavior but in fact, elicited active avoidance of the APP odor.

**Figure 7:**
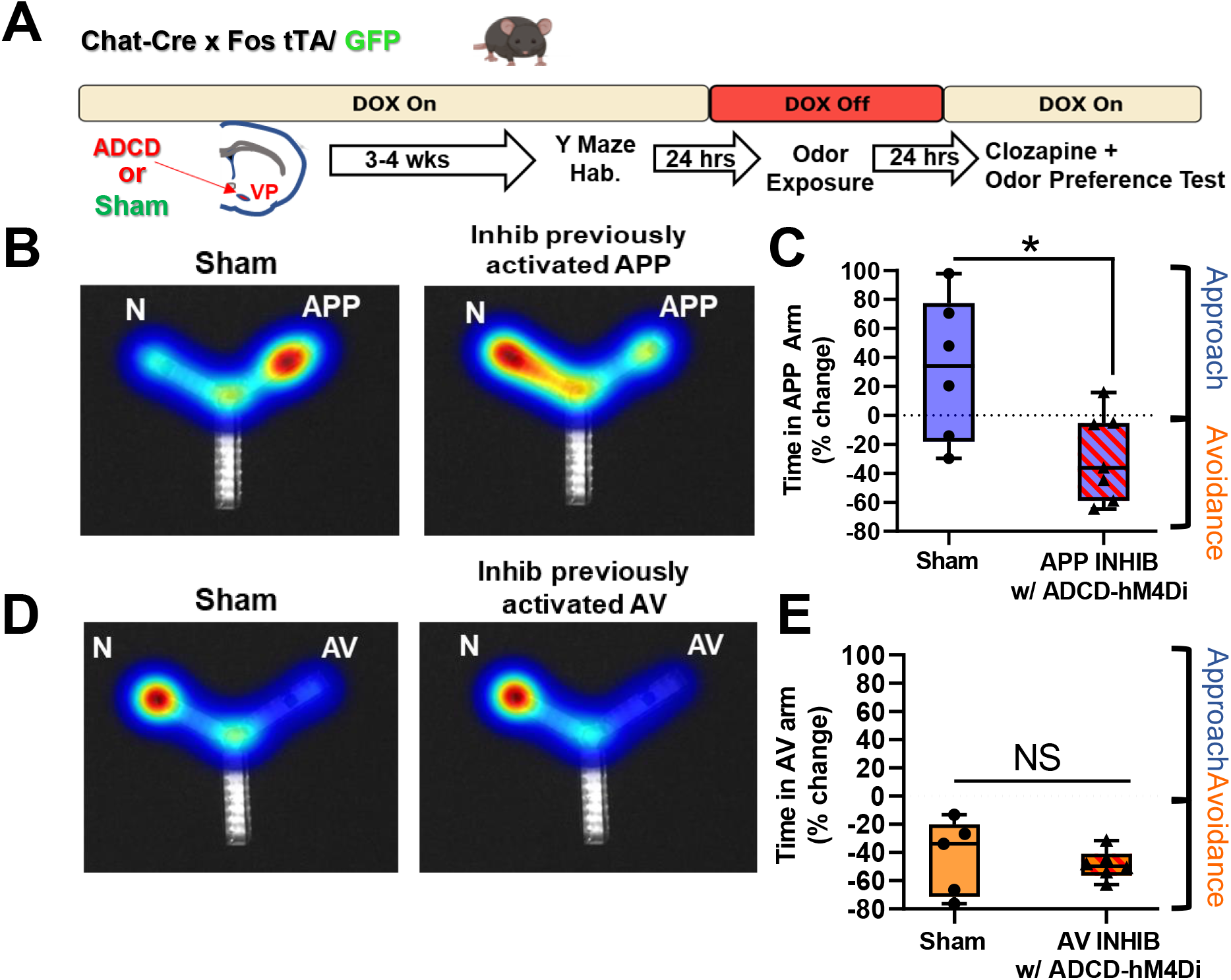
Selective chemogenetic inhibition of previously activated APP vs. AV cholinergic neurons in the VP abolishes approach to the appetitive odor, reversing behavior to strong avoidance. **A.** Workflow and timeline of behavior experiments using ADCD to target inhibitory DREADD’s for specific inhibition of previously activated subpopulations of VP cholinergic neurons. Chat-Cre x Fos-tTA/GFP mice were injected with ADCD and AAV-Syn-eGFP mice in the VP. Mice in the sham group were only injected with AAV-Syn-GFP but otherwise went through identical procedures as ADCD injected mice. Following recovery from surgery, mice were habituated (2 x 10 min) in the Y-Maze. Following arena habituation, mice were removed from a DOX diet, thus allowing for ADCD expression. Approximately 24-hours later, mice were exposed to either the appetitive odor (APP) or aversive odor (AV) in one arm of the Y-Maze, thus labeling activated VP cholinergic neurons with ADCD. Following odor exposure, mice were put back on a DOX diet, preventing further ADCD expression. The ADCD construct labels activated neurons with an inhibitory DREADD (ADCD-hM4Di). Approximately 24-hours later, all mice were injected IP with 0.1 mg clozapine to inhibit VP cholinergic neurons that were previously activated by the APP or AV odor . Approximately 15-minutes later, mice underwent an odor preference test for the odor to which they had been previously exposed. **B.** ADCD-hM4Di mediated inhibition of VP cholinergic neurons previously activated in response to the appetitive odor (APP) not only blocks approach behavior, but reverses the approach behavioral response to active avoidance. Left: representative heatmap showing approach behavior to the APP odor in a sham mouse (injected with AAV-Syn-eGFP). Right: Representative heatmap in a mouse following inhibition of VP cholinergic neurons previously activated in response to the APP odor, now displaying active avoidance behavior in response to the APP odor (i.e., more time spent in the N arm). **C.** Mice in the sham group (n = 6) exhibit approach to the APP odor. Inhibiting previously activated VP cholinergic neurons in response to APP with ADCD-hM4Di + 0.1 mg/kg clozapine (n = 7), leads to significantly more time in the saline paired arm, consistent with both a decrease in approach behavior and reversal of approach behavior to active avoidance. * *p* < 0.05. **D.** ADCD-hM4Di mediate inhibition of VP cholinergic neurons previously activated in response to the aversive odor (AV). Left: Representative heatmap depicting avoidance of the AV odor in a control mouse. Right: ADCD inhibition of VP cholinergic neurons previously activated in response to the AV odor has no effect on innate avoidance of the AV odor and mice continue to exhibit avoidance behavior. **E.** Mice in the sham group (n = 5) exhibit innate avoidance of the AV odor. Mice with ADCD-hM4Di + 0.1 mg/kg clozapine (n = 6) also exhibit avoidance of the AV odor. (* *p* < 0.05).

In a preference test with the AV odor, mice in the sham group displayed avoidance of the AV odor (Fig 7D left and 7E). In sharp contrast to the dramatic effects of silencing APP odor activated VP cholinergic neurons, mice with ADCD-hM4Di silencing of VP cholinergic neurons previously activated by exposure to the AV odor had no effect: mice continued to display avoidance of the AV odor (Fig 7D right and 7E). These results underscore our findings using hM4Di for non-selective inhibition of VP cholinergic neurons (Fig 4) and were devoid of any changes in locomotor activity (Fig S7). Combined, our DREADD inhibition experiments show that inhibition of VP cholinergic neurons (either general silencing or inhibiting APP VP cholinergic neurons) reversed approach behavior and led to active avoidance of the APP odor.

### APP vs. AV odor activated VP cholinergic neurons are intermingled, but differ in certain aspects of electrophysiological properties, neuronal morphology, and projections to the BLA

We next examined features that might be shared and factors that might distinguish the two subpopulations of VP cholinergic neurons. First, for all mice assessed in Figures 5 and 6 above, we relocalized the site of ADCD injection and mapped APP odor responding (Fig 8, blue circles) and AV odor responding (Fig 8, orange triangle) cholinergic neurons. Across the targeted regions (bregma +0.62 to +0.14) we found that these differentially activated subgroups of VP cholinergic neurons are spatially intermingled within the VP (Fig 8). Based on these findings, there does not seem to be a spatial segregation of APP vs. AV odor activated VP cholinergic neurons.

**Figure 8:**
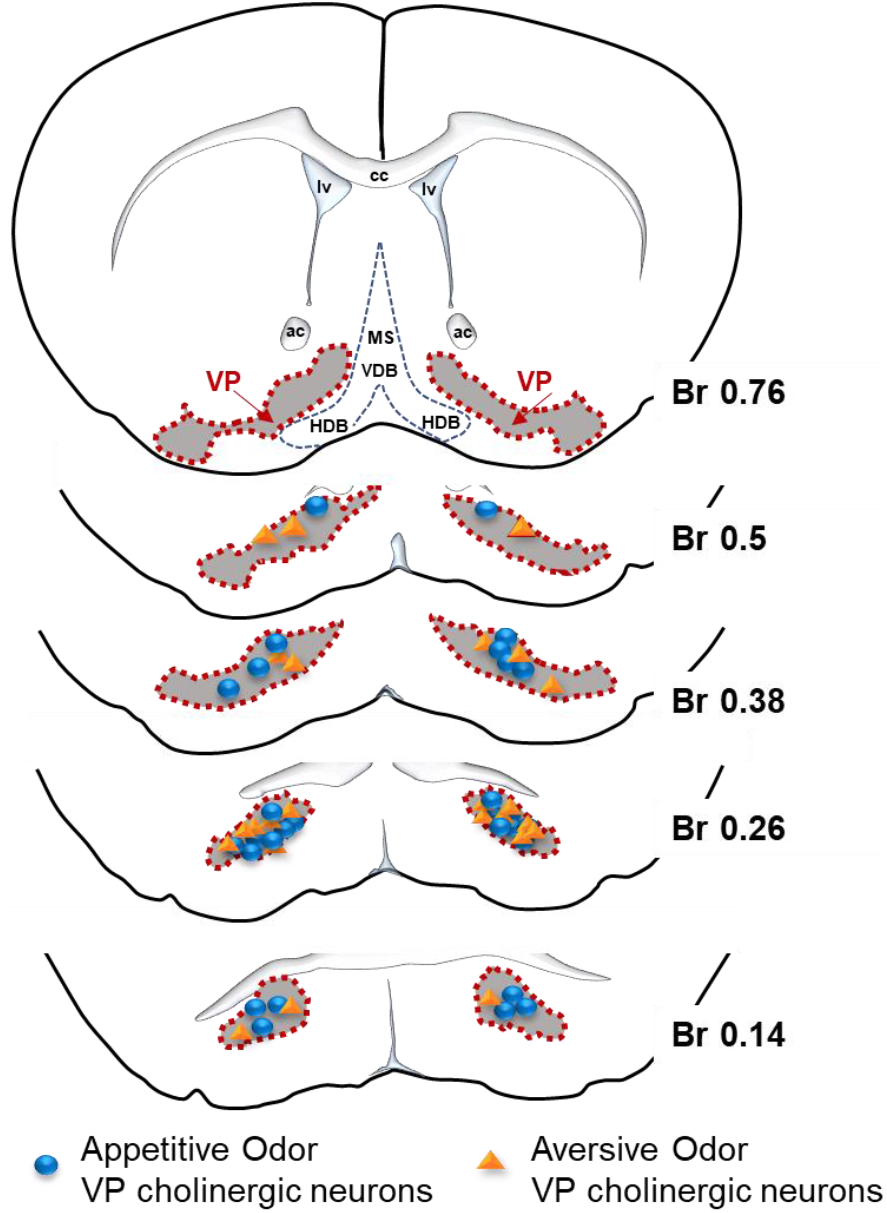
Appetitive and aversive odor activated VP cholinergic neurons are intermingled within the VP. Viral injection sites from mice studied in Figures 5 & 6 were used to relocalize the approximate positions of VP cholinergic neurons activated by the appetitive odor vs. those activated by the aversive odor. Appetitive and aversive odor activated VP cholinergic are intermingled across the anterior-posterior axis of the VP assayed (Bregma +0.76 – Bregma +0.14).

Next, to ask how the two subpopulations of VP cholinergic neurons might differ in electrophysiological properties from each other, we used a Cre-dependent activity marker (FLEX-RAM; (Sørensen et al., 2016)) to label each population as described above with ADCD, and then used patch-clamp recordings to quantify 18 different aspects of the electrophysiological profile of each subpopulation (Fig 9A and 9B). Recordings were performed at least 3 days after odor exposure, to avoid any transient changes in functional profile that might arise due to acute neuronal activation. We compared the properties of APP odor activated and AV odor activated VP cholinergic neurons to a pool of “non-activated” VP cholinergic neurons obtained from Chat-tau-eGFP mice maintained in the home cage. Most passive and active features were shared by VP APP and AV cholinergic neurons (Fig S8). However, there were important differences between APP and AV odor activated VP cholinergic neurons that produced clear distinctions in their action potential profiles and their relative excitability. Specifically, AV odor activated VP cholinergic neurons were slightly more hyperpolarized than APP activated VP cholinergic neurons (Fig 9C and 9F). AV odor activated VP cholinergic neurons also had a shorter latency to firing (Fig 9E and 9G) and had a smaller amplitude and shorter duration of the afterhyperpolarization potential (Fig 9D and 9H). The differences in these three features are detailed in Fig 9 (Fig 9F – 9H). The net effect of the differences between these three features underlie the phenotype that APP VP cholinergic neurons are less excitable and less prone to repeat firing than AV VP cholinergic neurons.

**Figure 9:**
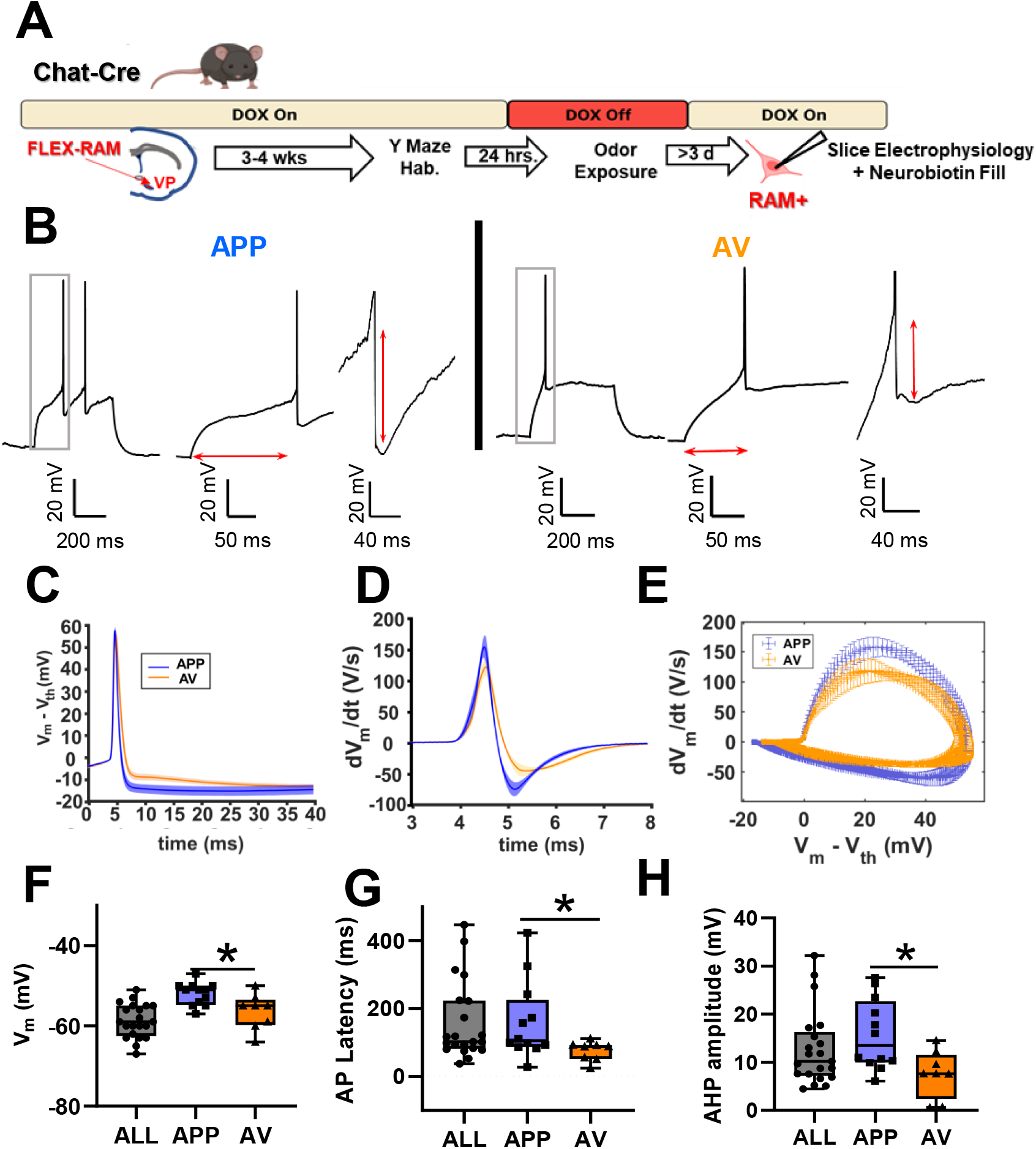
Differences in electrophysiological properties between appetitive odor activated vs. aversive odor activated VP cholinergic neurons. **A.** Timeline of slice electrophysiology experiments using AAV.RAM.d2TTA::TRE.FLEX.tdTomato (FLEX-RAM) labeling of activated VP cholinergic neurons. Chat-Cre mice were injected with FLEX-RAM in the VP. Following recovery from surgery (3 - 4 weeks) mice were habituated in the Y-Maze (2 x 10 min). Following arena habituation, DOX diet was removed and replaced with regular chow to allow FLEX-RAM expression. After 24 hours, mice were exposed to either the appetitive odor (APP) or aversive odor (AV) in one arm of the Y-Maze with the other arm blocked, thus labeling activated VP cholinergic neurons with FLEX-RAM. Mice were then returned to a DOX diet to block further expression of FLEX-RAM. Following a minimum of 3 days (to avoid assessing transient changes in electrophysiological properties due to changes in immediate early genes), coronal slices containing the VP were taken for slice electrophysiology. During slice electrophysiology recordings, the patch pipette was filled with neurobiotin for subsequent relocalization and reconstruction of FLEX-RAM labeled neurons (Fig 9). **B.** Representative electrophysiology traces from an appetitive odor activated VP cholinergic neuron (left, APP, n = 12) and an aversive odor activated VP cholinergic neuron (right, AV, n = 8). Left traces = representative traces at rheobase. Middle traces = rheobase traces zoomed in to show differences in latency to fire an action potential. Right traces = rheobase traces zoomed in to illustrate differences in amplitude of the hyperpolarization potential. **C.** Extended time course of action potential currents (V_m_ – V_th_) for APP vs. AV VP cholinergic neurons reveals the deeper and more prolonged nature of the AHP in APP VP cholinergic neurons. **D.** Action currents of APP vs. AV VP cholinergic neurons, plotted as dV/dt vs. time, highlighting visualization of differences in short term AHP. **E.** Phase plots of action potential dynamics (dV/dt vs. V) providing visualization of multiple features of the waveform including spike threshold (far left), upstroke velocity, spike amplitude (far right), downstroke velocity and afterhyperpolarization to return to Vm. **F - H.** Comparison of electrophysiological properties of identified APP vs. AV VP cholinergic neurons with VP cholinergic neurons from home-cage Chat-tau-eGFP mice not exposed to either odor. APP VP cholinergic neurons differ from AV VP cholinergic neurons in **(F)** resting membrane voltage, **(G)** AP latency, and **(H)** AHP amplitude. * *p* < 0.05.

All cells examined during electrophysiological recordings were identified by post-hoc ChAT IHC and by expression of FLEX-RAM. During patch clamp recordings, we also filled the neurons with neurobiotin and subsequently reconstructed the neuronal morphology of APP odor and AV odor activated VP cholinergic neurons (Fig 10 and 10B). A convex hull analysis was conducted to assess the surface area and volume occupied by APP vs. AV odor activated VP cholinergic neurons. The convex hull enveloping APP odor activated VP cholinergic neurons was significantly smaller in surface area (*t* (10) = 2.70, *p* < 0.05; Fig 10C) and volume (*t* (10) = 2.32, *p* < 0.05; Fig 10D) compared to AV odor activated VP cholinergic neurons. APP odor activated VP cholinergic neurons also exhibited a significantly smaller dendrite area (*t* (14) = 3.75, *p* < 0.05; Fig 10E) and dendrite volume (*t* (14) = 5.14, *p* < 0.05); Fig 10F). The maximum length of reconstructed segments was also significantly smaller in APP odor activated VP cholinergic neurons (*t* (14) = 3.49, *p* < 0.05; Fig 10G). In addition to these features, APP odor activated VP cholinergic neurons had a significantly greater number of intersecting points in a sholl analysis at distances close to the soma (at 10 µm *t* (14) = −2.69, *p* < 0.05; at 20 µm *t* (14) = −3.16, *p* < 0.05; Fig 10H). Next, we assessed the percentage of reconstructed neurons whose dendrites reached a set distance from the soma in a sholl analysis. The dendritic arbor of ∼ 80% of AV odor activated VP cholinergic neurons extended at least 100 µm from the soma, whereas less than 20% of APP odor activated VP cholinergic neurons reach this distance (Fig 10I). Additional morphometric features that were not statistically different between APP odor and AV odor activated VP cholinergics neurons are shown in Figure S9. Overall, these results consistently demonstrate APP odor activated VP cholinergic neurons are smaller and more complex in their proximal dendritic arbor than AV VP cholinergic neurons. These data, along with our slice electrophysiology profile of APP and AV odor activated VP cholinergic neurons, are consistent with potential differences in their circuit engagement.

**Figure 10:**
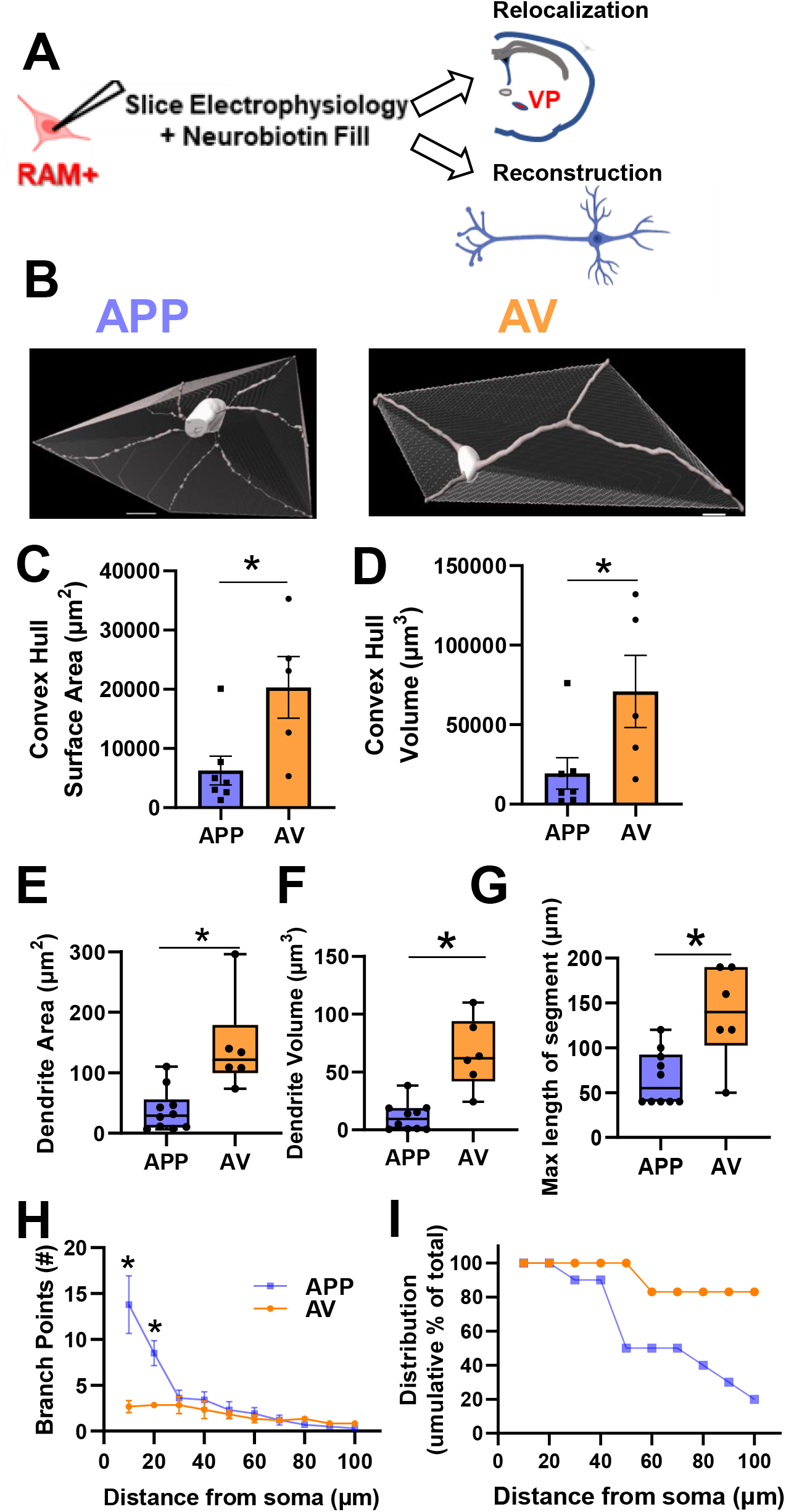
Differences in neuronal morphology between appetitive odor activated vs. aversive odor activated VP cholinergic neurons. **A.** FLEX-RAM labeled neurons were filled with neurobiotin during slice electrophysiology recordings (see Fig 8 for details). Following electrophysiology recordings, slices were post-fixed in 4% PFA for 24-hours and then 1x PBS until immunostaining. Slices were stained for ChAT (to visualize cholinergic neurons) and streptavidin (to visualize neurobiotin filled cells). Following immunostaining, slices were imaged on a confocal microscope to visualize FLEX-RAM, ChAT, and streptavidin. **B.** Confocal images were reconstructed using Imaris software. Reconstructed neurons from an appetitive odor activated (APP) VP cholinergic neuron (n = 10, left) and an aversive odor activated (AV) VP cholinergic neuron (n = 6, right), with a convex hull analysis. A convex hull is a 3D polyhedron that encompasses all distal points of the reconstructed neuron. Scale bar is 10 µm **C – D.** Surface area and volume measurements of the convex hull. The convex hull from APP VP cholinergic neurons are significantly smaller in **(C)** surface area and **(D)** volume. * *p* < 0.05 **E – G.** APP VP cholinergic neurons are smaller and have fewer linear dimension features vs AV VP cholinergic neurons. APP VP cholinergic neurons exhibit a significantly smaller **(E)** dendrite area, **(F)** dendrite volume, and **(G)** max length of reconstructed segment. * *p* < 0.05. **H.** Sholl analysis of reconstructed APP and AV VP cholinergic neurons. Despite exhibiting reduced morphometric properties, APP VP cholinergic neurons display greater proximal soma complexity in a sholl analysis. APP VP cholinergic neurons display a significantly greater number of branch points at 5 and 10 µm from the soma. * *p* < 0.05. **I.** Although AV VP cholinergic neurons display less branching vs. APP VP cholinergic neurons, AV neurons are overall lengthier. A larger percentage of AV VP cholinergic neurons are able to reach greater distances away from the soma vs. APP VP cholinergic neurons.

The BLA is a major projection target of VP cholinergic neurons (Root et al., 2015; Záborszky et al., 2018). To begin to examine how these two distinct populations of VP cholinergic neurons might differ in their projections encoding valence, we examined the relative predominance (or lack thereof) of the APP vs. AV projections to the BLA. We chose to specifically focus on the BLA since it receives known cholinergic input from the VP (Root et al., 2015; Záborszky et al., 2018). We assessed the relationship between BLA-projecting and activated cholinergic neurons in the VP following odor exposure (Fig 11B and 11C). Although consistent with our findings above that there were an equal number of VP cholinergic neurons that were activated by the APP vs. AV odor (Fig 11D left), the relative proportion of BLA-projecting APP vs. AV VP cholinergic neurons was not equivalent. The percentage of BLA-projecting VP cholinergic neurons that were activated by the AV odor was significantly greater than the number of BLA-projecting VP cholinergic neurons activated by the APP odor (*t* (6) = −2.98, *p* < 0.05; Fig 11D right). This result underscores the predominance of AV encoding VP cholinergic projections to the BLA.

**Figure 11:**
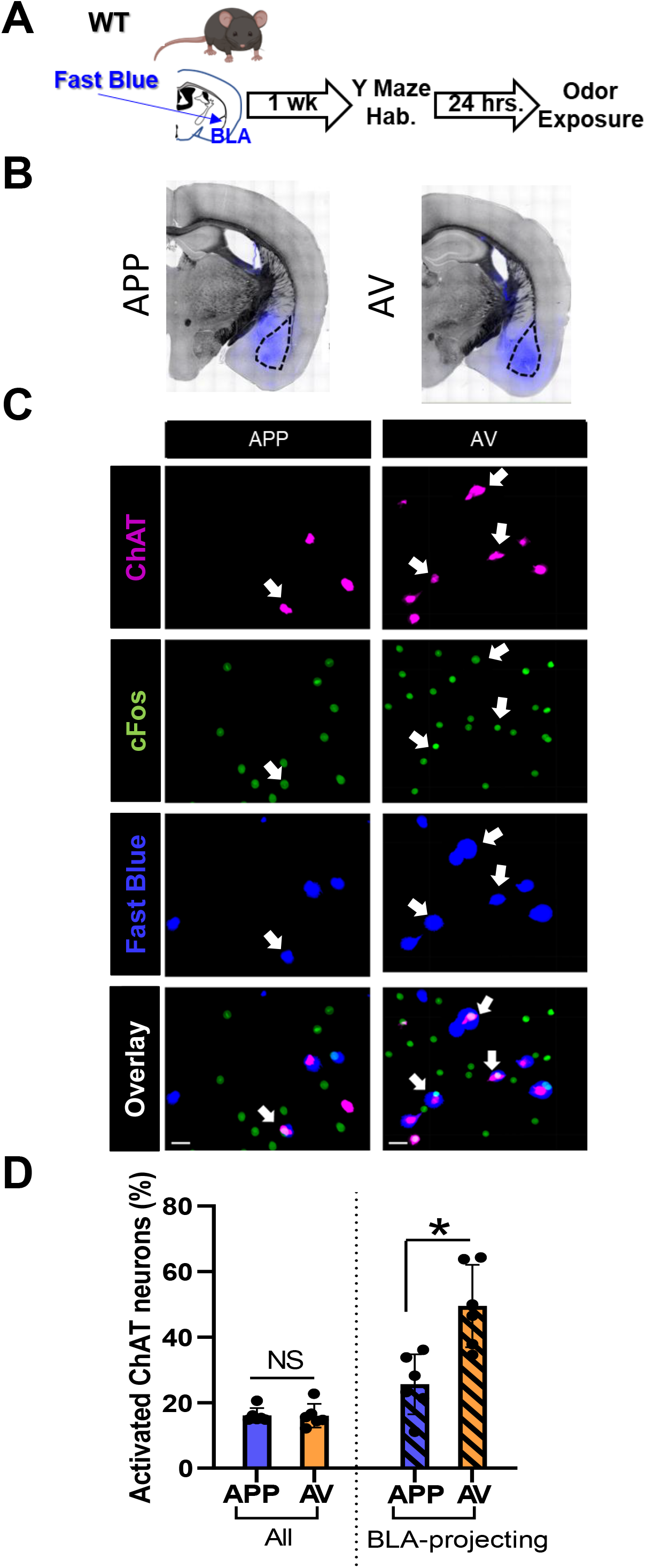
Although both APP and AV VP cholinergic neurons project to the BLA, the predominant BLA input from the VP stems from AV encoding VP cholinergic neurons. **A.** Workflow and timeline of experiment using Fast Blue to retrogradely label BLA-projecting VP cholinergic neurons. Following recovery from surgery (∼1 week), mice were habituated in the Y-Maze (2 x 10 min). Approximately 24-hours later, mice were exposed to either the appetitive odor (APP) or aversive odor (AV) in one arm of the Y-Maze. 45-minutes following odor exposure, mice were euthanized, and tissue was processed for ChAT and cFos IHC. **B.** Representative images in mice injected with Fast Blue in the BLA (in blue) and overlayed on a bright-field image. Mice were either exposed to the APP odor (left) or AV odor (right). **C.** Representative images from the VP in a mouse exposed to the APP odor (left column) and AV odor (right column). First row = ChAT (magenta, cholinergic marker), second row = cFos (green, marker of neuronal activation), third row = Fast Blue (blue, BLA-projecting VP neurons), bottom row = overlay of all channels. Scale bar is 20 µm. **D.** There is no difference between APP vs. AV odor exposed mice in terms of the overall percentage of odor activated VP cholinergic neurons (left). However, the AV responsive VP cholinergic neurons constitute a significantly greater percentage of BLA-projecting VP cholinergic neurons. * *p* < 0.05.

## Discussion

We identified two non-overlapping subpopulations of VP cholinergic neurons-one activated in response to an APP odor, and a second, distinct subpopulation activated in response to an AV odor. We demonstrate that despite being activated (both in terms of an increase in cFos expression and an increase in calcium activity) in response to both the APP odor and AV odor, VP cholinergic neuron activity is absolutely required for approach behavior. These two subpopulations of VP cholinergic neurons also differ in passive and active electrophysiological properties, dendritic morphology, and projections to the BLA. These data are consistent with differential integration of each subpopulation into valence processing circuits.

### VP cholinergic neurons are engaged in behavioral responses to both the APP and AV odor

Very few studies have examined how mice respond to innately pleasant stimuli. The APP odor used in the present study was 2-phenylethanol, the active component of rose oil (Ueno et al., 2019). A previous experiment showed that although mice do not exhibit a preference for 2-phenylethanol, mice display anti-depressive like behaviors (Ueno et al., 2019). We demonstrate here that mice display approach to the APP odor. One hypothesis for this difference is the arena in which the mice were tested. We used two arms of a Y-maze in which mice were given the ability to freely roam between two arms, likely leading to the exhibition of approach behavior. Although the VP has a well-established role in reward-related behaviors such as drug-addiction (Smith et al., 2009; Morales and Berridge, 2020; Ottenheimer et al., 2020), the role of VP cholinergic neurons in reward behaviors is unclear. A previous experiment found that 192-IgG-Saporin induced lesion of cholinergic neurons in parts of the NAc and VP reduced cocaine self-administration (Smith et al., 2004), implicating VP cholinergic neurons in some aspect of reward related behaviors. Our results show that VP cholinergic neurons are engaged following exposure to an APP stimulus.

Mice reliably display innate avoidance behavior following predator odor exposure (Pérez-Gómez et al., 2015; Rosen et al., 2015; Hwa et al., 2019; Wu et al., 2019; Barbano et al., 2020). This behavior is consistent across multiple predator odors (fox urine, cat urine, mountain lion urine, or 2,5-Dihydro-2,4,5-trimethylthiazoline (TMT)). Avoidance of the predator odor is accompanied by numerous neurobiological changes. Amongst a wide array of effects, the most notable include: cFos activation in brain regions related to stress (Janitzky et al., 2015), the medial prefrontal cortex (mPFC) (Hwa et al., 2019), and the BLA (Butler et al., 2011). The results from the current study add to these findings and along with our previous work (Rajebhosale et al., 2021), demonstrate that activation of VP cholinergic neurons is another reliable response to predator odor exposure.

### VP cholinergic signaling is required for approach behavior

While VP cholinergic neurons were activated in response to both the APP and AV odor, silencing VP cholinergic neurons (either general inhibition or silencing of previously activated APP subpopulation) resulted in changes to approach but not avoidance behaviors. Inhibition of VP cholinergic neurons eliminated approach to the APP odor. In fact, these mice displayed behaviors consistent with avoidance of the APP odor.

Numerous studies have reported that approach behaviors are more readily manipulated than avoidance behaviors (Douton et al., 2021; Gil-Lievana et al., 2022; Yu et al., 2022). Our results show general silencing of VP cholinergic neurons, as well as inhibiting APP odor activated VP cholinergic neurons, not only blocks approach behavior, but reversed the behavior leading to avoidance of the APP odor. This suggests avoidance may be the default behavior until mice encounter an APP stimulus which engages APP VP cholinergic neurons. BFCN’s outside of the VP have also been shown to play a role in behavioral responses to an APP stimulus (Teles-Grilo Ruivo et al., 2017; Borden et al., 2020; Crouse et al., 2020). However, these previous studies focused on learned behaviors associated with rewarding stimuli, as opposed to innate behaviors. We have previously found that distinct subsets of BFCN’s govern learned vs. innate behavioral responses (Rajebhosale et al., 2021). As such VP cholinergic neurons may play a unique role in innate behavioral responses to salient stimuli. While avoidance behaviors are governed by multiple circuits (discussed below) and thus resistant to inhibition of a single component, approach behaviors are more susceptible to inhibition of VP cholinergic neurons. Furthermore, local cholinergic signaling within the VP may influence behavioral outcomes. A recent experiment highlighted the importance of local cholinergic signaling within the VP in regulating pain perception (Ji et al., 2023). Therefore, inhibiting APP odor activated VP cholinergic neurons may in turn activate AV odor VP cholinergic neurons, thus leading to avoidance of the APP odor.

Avoiding predators and noxious stimuli is crucial for survival. As such, the numerous brain regions and circuits known to be involved in avoidance of the predator odor (Hebb et al., 2004; Butler et al., 2011; Janitzky et al., 2015; Pérez-Gómez et al., 2015; Rosen et al., 2015; Hwa et al., 2019; Wu et al., 2019; Barbano et al., 2020) might provide redundancy and compensate for the inhibition of AV odor activated VP cholinergic neurons, thus preserving a behavior critical for survival. In addition, cholinergic signaling from other basal forebrain cholinergic neurons (BFCN’s) also contribute to behavioral responses to AV stimuli. Cholinergic projections from the nbM to BLA are critical for cue associated fear learning (Jiang et al., 2016; Knox, 2016; Rajebhosale et al., 2021; Crimmins et al., 2022), and projections from the MS/DBB to the hippocampus are associated with stress and anxiety (Eck et al., 2020; Mineur and Picciotto, 2021; Ren et al., 2022). Cholinergic signaling outside of the basal forebrain also plays a role in avoidance behaviors (Aitta-Aho et al., 2018). These compensating brain regions and circuits (both cholinergic and non-cholinergic) may allow an animal to maintain continued avoidance of threatening stimuli, despite the inhibition of VP cholinergic neurons. In contrast, our results demonstrated that VP cholinergic signaling is required for behavioral responses towards a rewarding stimulus. Inhibition of VP cholinergic neurons not only abolished approach behavior but led to active avoidance of an APP stimulus.

### Functional heterogeneity of VP cholinergic neurons

Our results demonstrate two distinct subpopulations of VP cholinergic neurons that are differentially activated by APP vs. AV odors. These two subpopulations of VP cholinergic neurons shared similarities in cFos expression, calcium signaling, anatomical location and most electrophysiological properties. Our experiments also identified significant differences between APP and AV odor activated VP cholinergic neurons. These differences further support the notion of the presence of two distinct subpopulations of VP cholinergic neurons.

In addition to differences in behavioral responses to VP cholinergic inhibition (described above), APP vs. AV VP cholinergic neurons exhibited differences in some electrophysiological features, morphology, and projections to the BLA. Notably, AV VP cholinergic neurons were more excitable than APP VP cholinergic neurons. AV VP cholinergic neurons displayed larger and longer dendritic arbors, but less complexity in their proximal dendritic arbors. Changes in neuronal morphology and alterations in electrophysiological features are two common indicators of changes in the connectivity of neural circuits (Zhu et al., 2016; Peng et al., 2021; Ciganok-Hückels et al., 2022; Udvary et al., 2022). As the primary output of VP cholinergic neurons targets the BLA (Root et al., 2015; Záborszky et al., 2018), we chose to examine the proportion of BLA-projecting VP cholinergic neurons activated by each odor. Our retrograde mapping experiment revealed that although both subpopulations of VP cholinergic neurons do indeed project to the BLA, a greater percentage of BLA-projecting VP cholinergic neurons were activated by the AV odor. Collectively, these results illustrate the presence of two distinct, non-overlapping subpopulations of cholinergic neurons in the VP, demonstrating functional heterogeneity of VP cholinergic neurons.

VP cholinergic neurons, along with cholinergic neurons found within the medial septum/diagonal band of Broca (MS/DBB), horizontal limb of the diagonal band (hDB), and nucleus basalis of Meynert (nbM) constitute cholinergic neurons located in the basal forebrain (BFCN’s). Emerging evidence illustrates functional heterogeneity of BFCN’s (Ananth et al., 2023). For example, cholinergic projections from the nbM to the BLA play a role in both cue-associated fear learning (Jiang et al., 2016; Knox, 2016; Wilson and Fadel, 2017; Rajebhosale et al., 2021; Crimmins et al., 2022) and reward learning (Aitta-Aho et al., 2018; Crouse et al., 2020). Cholinergic projections from the MS/DBB to the hippocampus have been heavily implicated in behavioral response to stress and anxiety, as well as in depressive-like behaviors (Paul et al., 2015; Mei et al., 2020; Mineur and Picciotto, 2021; Mineur et al., 2022). Recent studies indicate MS/DBB projections to the hippocampus also play a role in reward learning (Teles-Grilo Ruivo et al., 2017; Borden et al., 2020). These studies show BFCN’s have an active role in encoding responses to both negative and positive valence stimuli. This is supported by in-vivo recordings of BFCN’s, which demonstrate increased firing in response to both reward and punishment (Hangya et al., 2015; Laszlovszky et al., 2020). These studies illustrate the incredible functional diversity and heterogeneity of BFCN’s. BFCN’s in one brain region can mediate behavioral responses to both positive and negative valence stimuli.

Despite the well-established role of the VP in valence processing, prior studies have not tested the participation of VP cholinergic neurons in valence encoding. Our results begin to fill this gap by demonstrating that VP cholinergic neurons are indeed, activated in response to both APP and AV odors. Moreover, our results show distinct and non-overlapping subsets of cholinergic neurons in the VP are activated in response to APP vs. AV odors. However, despite the activation of VP cholinergic neurons in response to both APP and AV odors, silencing VP cholinergic neurons (general inhibition or inhibiting previously activated subpopulations) only affected approach behavior. These differential behavioral effects following inhibition of VP cholinergic neurons further lends support to our finding of two distinct subpopulations of VP cholinergic neurons.

## Conclusions

The results from this study demonstrate two distinct subpopulations of cholinergic neurons exist in the VP which differentially encode valence of olfactory stimuli. The two subpopulations of VP cholinergic neurons differ in certain electrophysiological properties, neuronal morphology, projections to the BLA, and behavioral responses following inhibition. Notably, general silencing of VP cholinergic neurons or inhibiting previously activated APP VP cholinergic neurons reversed approach behavior and led to the display of avoidance behavior. These results reveal circuit-level differences between the two subpopulations of VP cholinergic neurons.

This present study focused specifically on the functional heterogeneity of VP cholinergic neurons. However, we cannot disregard cholinergic signaling within other parts of the basal forebrain and other brain regions that contribute to valence encoding. Numerous studies have reported the activation of BFCN’s following exposure to both rewarding and aversive stimuli. Although VP cholinergic neurons were hypothesized to resemble other BFCN’s and encode salience, we show that cholinergic projection neurons of the VP are functionally complex and demonstrate the capability to encode valence of olfactory stimuli. In order to better determine the role of VP cholinergic neurons in encoding valence, future research can determine if these results extend to valence of other sensory stimuli (i.e., taste, touch), and to learned (i.e., conditioned) responses. Finally, we began to examine how these two subpopulations of VP cholinergic neurons differ by examining one specific projection to the BLA. Future studies can examine potential differences in other brain regions that receive cholinergic input from the VP (mPFC and MD). Moreover, another area for future research is to determine how valence encoding VP cholinergic neurons interact with valence encoding neurons of the BLA and/or other regions, to promote approach vs. avoidance behaviors. Targeting positive vs. negative valence encoding microcircuits may lead to the development of more efficacious pharmacotherapeutic treatments for neuropsychiatric disorders which are characterized by misattributions of valence and/or motivational imbalance.

This work was supported by funds from the National Institute of Neurological Disorders and Stroke (NINDS) Intramural Research Program (IRP) (1ZIANS009424 to David A. Talmage, 1ZIANS009416 and 1ZIANS009422 to Lorna W. Role).

## Author Contributions

R.K., M.A., L.W.R., and D.A.T. designed the experiments. R.K., M.A., and N.S.D. performed the experiments and analyzed the data. R.K., L.W.R., and D.A.T. wrote the article.

## Methods

### Animals/Subjects

8-12 week old male and female mice were randomly assigned to groups at the start of all experimental procedures. The strains of mice used are: C57BL/6J (Jax #000664), Chat-IRES-Cre::Δneo (B6; 129S6-Chattm2(cre)Lowl/J; Jax # 006410, abbreviated Chat-Cre), and the offspring of Chat-Cre mice crossed to Fos-tTA/Fos-shEGFP mice, (B6. Cg-Tg(Fos-tTA, Fos-EGFP*)1Mmay/J; Jax # 018306, abbreviated Chat-Cre x Fos-tTA/GFP). Chat-tau-eGFP mice (a gift from S.Vijayaraghavan, University of Colorado (Grybko et al., 2011)), which express GFP in all cholinergic neurons and processes, were used as naïve home-cage controls in electrophysiology experiments. Previous studies indicate the electrophysiological profile of neither Chat-tau-eGFP or Chat-Cre mice differ from non-genetically tagged cholinergic neurons (López-Hernández et al., 2017). All mice were housed in a 12-hour light/dark cycle with temperature and humidity control. Food and water were available ad-libitum. All proceudres were approved by the NINDS Animal Care and Use Committee (ASP # 1531). For all experiments, the APP odor used was 2-phenylethanol (Sigma # 77861), and the AV odor used was mountain lion urine (Predator Pee # 92016).

### Surgery

Mice were anesthetized using isoflurane (3-5% induction, 1.5% maintenance) and placed in a stereotax. Bilateral injections targeted the VP using coordinates from the Paxinos Mouse Brain Atlas (AP + 0.38, ML +/- 1.45, DV 5.0). A Hamilton syringe was slowly lowered to the VP and mice were injected with ∼0.3 µl of the following viral vectors dependent on experiment: AAV_9_.DIO.TRE.hM4Di.P2A.mCherry (ADCD; (Rajebhosale et al., 2021) packaged at UNC viral vector core), AAV_9._RAM.d2tTA.TRE.mCherry.NLS-FLAG (RAM; (Sørensen et al., 2016); Addgene # 63931, packaged at UNC viral vector core), AAV_9._Syn.Flex-GCaMP6f.WPRE.SV40 (GCaMP6F; Addgene # 100837), AAV_8_.hSyn.DIO.hM4Di (Gi).mCherry (hM4Di; Addgene # 44362), AAV_9_.eSyn.eGFP (Vector Biolabs # VB4870), AAV_9._RAM.d2tTA.TRE.Flex.tdTomato (FLEX-RAM; Addgene # 84468, packaged at NINDS Viral Vector Core). For retrograde tracing experiments, 0.15 µl Fast Blue (Polysciences # 17740-1) was injected bilaterally in the basolateral amygdala (BLA, AP −1.35, ML +/- 3.2, DV 4.7). For optical recording of calcium signaling, in-vivo fiber photometry was used. A fiber optic cannula (Neurophotometrics, ferrule diameter 1.25 mm, core diameter 400 µm) was implanted ∼100 µm above the VP virus injection site and secured using dental cement.

### Behavior

#### Odor Preference Behavior Tests

All experiments assessing the behavioral responses to odors were conducted using a near-infrared (NIR) modular Y-Maze with a NIR camera (Med Associates). Initial odor preference was assessed in a single 10-minute session. One arm of the Y-Maze contained an odor (either APP or AV) on a gauze pad and one arm contained saline (null odor). Mice began the session in the third vacant arm. After initial entry into either the odor-containing or saline-containing arm, re-entry into the starting arm was blocked, forcing mice to choose between the odor vs. saline arm. Time spent in each arm (odor vs. saline) was analyzed using Ethovision software.

#### ADCD/RAM labeling of activated VP cholinergic neurons

For ADCD and RAM labeling experiments, Chat-Cre x Fos-tTA/GFP mice were given approximately 3-4 weeks to recover from surgery post viral injection. Following recovery, behavior sessions occurred over three consecutive days. On the first day (Day 0), mice were habituated twice to the Y-Maze. During each habituation session, mice were allowed to explore only one arm of the Y-Maze for 10 minutes, counter-balanced for each arm of the Y-Maze. Following the second habituation session, mice were transferred to a clean cage and removed from a doxycycline (DOX) diet, thus allowing for ADCD or RAM expression. On Day 1, approximately 24 hours later, mice were exposed to either an odor or saline in one arm of the Y-maze (labeling activated VP cholinergic neurons). Following ADCD or RAM labeling of activated VP neurons, mice were returned to a DOX diet. On Day 2, approximately 24 hours later, mice were then exposed to either saline or an odor in a different arm of the Y-Maze. Mice were returned to the home cage until tissue was collected for additional processing.

#### DREADD inhibition experiments

Odor preference was assessed following general inhibition of VP cholinergic neurons using a floxed inhibitory DREADD in Chat-Cre mice (hM4Di), or inhibition of previously activated VP cholinergic neurons (ADCD-hM4Di) in Chat-Cre x Fos-tTA/GFP mice. For hM4Di odor preference tests, mice were allowed to recover from surgery for 3-4 weeks and then injected IP with 0.1 mg/kg clozapine. Approximately 15 minutes later, odor preference was assessed in the Y-Maze, identical to the odor preference test described above. For ADCD labeling in odor preference tests, mice were allowed to habituate to the Y-Maze on Day 0 and activated VP

cholinergic neurons were labeled with ADCD on Day 1. For selective inhibition of previously activated VP cholinergic neurons on Day 2, mice were injected with 0.1 mg/kg clozapine and allowed to explore two arms of the Y-Maze (previously exposed odor vs. saline). Time spent in each arm (odor vs. saline) was analyzed using Ethovision software.

#### Analysis

All odor preference behavior videos were first converted to .mP4 files and uploaded to Ethovision (v.15 from Noldus). The Y-Maze dimensions were used to delineate the arena borders and each arm containing an odor or saline gauze pad was outlined as a zone. Automated rodent tracking was used for each video and the time (in seconds) the mouse spent in each zone was obtained.

### In-vivo calcium signaling assays using fiber photometry

#### Acquisition

Chat-cre mice were injected with a cre-dependent GCaMP6F in the VP and a fiber optic cannula was implanted above the viral injection site. Fiber photometry recordings were made using the Neurophotometrics FP3002 system. Acquisition parameters were: 415 (for isobestic channel recordings) and 470 (GCaMP channel recordings) nm channels, 20 frames/sec, power ∼100 µw. All fiber photometry recordings were conducted in behavior chambers equipped with an odor delivery system (Med Associates). Mice were given two days to habituate to the behavior chamber and patch cords. Mice were then exposed to timed delivery of either odor (APP or AV). Timed odor delivery took place in a 10-minute session, where odor was delivered (3 x 10 sec) every 3 minutes. Approximately 24 hours following the first odor exposure session, mice were exposed to the opposite odor.

#### Analysis

Processing of all fiber photometry data was performed using a custom MATLAB script. Initialization frames (∼5 sec at the start of recordings) were removed from the raw data and raw fluorescence values were binned at 5 Hz intervals. The GCaMP signal was then normalized to the isobestic signal and fluorescence measures were obtained. The change in fluorescence (ΔF/F) was calculated and converted to z-scores. The integrated z-score vs. time was estimated by the area under the curve (AUC), calculated using GraphPad Prism representing a single value combining fluorescence intensity and time. The max z-score ΔF/F was defined as the highest z-score ΔF/F during the 10 second odor exposure.

### Immunohistochemistry (IHC)

#### Tissue processing

Following behavior testing or odor exposure (∼45 minutes for C57 and Chat-Cre mice; ∼2.5 hours for Chat-Cre x Fos tTA/GFP mice), mice were perfused with phosphate buffered saline (PBS) and 4% paraformaldehyde (PFA). Brains were post-fixed in 4% PFA overnight before being cryoprotected in 30% sucrose. 50 µm slices were cut on a cryostat and stored in 50% PBS/glycerol at −20 °C until immunostaining.

#### Immunostaining

For all experiments, free floating slices were first washed (3 x 10 min) in PBS + 2% Triton X-100 (PBST). Slices were then transferred to a blocking solution containing PBST and 10% normal donkey serum (NDS) and incubated on a shaker for 2 hours at room temperature. Following blocking, slices were incubated overnight at 4°C in blocking solution with the addition of the following antibodies (dependent on experiment): goat anti-ChAT (Millipore Sigma AB144P), rabbit anti-cFos (Synaptic System 226 008), chicken anti-GFP (Abcam ab13970), mouse anti-mCherry (Living Colors 632543), or rat anti-Substance P (Millipore Sigma MAB356). All primary antibodies were used at 1:500. Following incubation with primary antibodies, slices were washed (3 x 10 min) in PBST and then incubated (2 hours at room temperature) in blocking solution with the addition of the following secondary antibodies (dependent on primary antibody used, all secondary antibodies were used at 1:1000): donkey anti-goat A647 (Invitrogen A-21447), donkey anti-rabbit A488 (Invitrogen A-21206), donkey anti-chicken A488 (Jackson Immuno 703-454-155), donkey anti-mouse A 594 (Invitrogen A-21203) or donkey anti-rat A 594 Invitrogen A-21209). Following incubation with secondary antibodies, slices were first washed in PBST (3 x 10 min) and then washed in PBS (1 x 10 min). Slices were mounted onto slides using Fluoromount-G with DAPI (Southern BioTech 0100-01).

#### Imaging

All images were acquired on an Olympus VS200 slide scanner with a Hamamatsu Orca-Fusion camera. Whole slice images were obtained using the 415, 470, 594 and 647 nm channels. Each image was acquired using 5 µm z-steps and identical exposure times for each channel. Images were saved as .vsi files before being converted to Imaris files for analysis.

#### Analysis

A pipeline for image analysis using Imaris is depicted in Figure S2. To assist in defining VP boundaries, immunostaining for Substance P, which demarcates VP borders (Root et al., 2015; Faget et al., 2018) was conducted in brain slices from a Chat-tau-eGFP mouse (Figure S3A). These boundaries were then used in later slices to delineate the VP. Whole slice images were first cropped and borders delineating the VP were created to mask any signal outside of the VP. The VP was identified as the brain region containing a sparse cluster of cholinergic neurons ventral to the anterior commissure and striatum, and dorsal to the densely packed cluster of cholinergic neurons in horizontal limb of the diagonal band. Within Imaris, the spots and surfaces function were used to create size and/or intensity-based thresholds for each channel. The colocalization tab was used to create a new channel with the colocalized signal between two defined channels. All data containing the total number of spots or surfaces was then exported.

### Electrophysiology

#### Odor Exposure

To assess passive and active electrophysiological properties of VP cholinergic neurons that had previously been activated in response to the APP or AV odor, we first labeled odor responsive neurons with the permanent activity marker FLEX-RAM (Addgene # 84468, (Sørensen et al., 2016)) in-vivo. FLEX-RAM is similar to RAM (i.e., activity dependent expression only when off a DOX diet) but is only expressed neurons that contain cre-recombinase. Therefore, Chat-Cre mice were injected with FLEX-RAM in the VP and exposed to either the APP or AV odor 3 - 4 weeks later (identical to the ADCD and RAM labeling experiments described above). Following odor exposure, mice were returned to a DOX diet and stayed in the home cage until electrophysiology recordings were conducted (a minimum of 72 hours). Odor naïve mice of Chat-tau-eGFP genetic background were used as non-odor exposed controls. Following a minimum of 72 hours, mice were euthanized, the brain was removed, and 300 µm slices containing the VP were taken on a Leica VT1200 vibratome (details discussed below).

#### Slice Preparation

Brain slices containing the VP were prepared using standard procedures (López-Hernández et al., 2017; Ting et al., 2018). Shortly after receiving a lethal dose of ketamine/xylazine, mice were perfused transcardially with an ice-cold cutting solution containing (in mM): 92 N-methyl-D-glucamine, 2.5 KCl, 1.25 NaH_2_PO_4_, 30 NaHCO_3_, 20 HEPES, 25 D-glucose, 2 thiourea, 5 ascorbate, 3 pyruvate, 0.5 CaCl_2_, and 10 MgSO_4_. Brains were removed as rapidly as possible, and a Leica VT1200 vibratome was used to make 300-µm-thick coronal sections. Slices were cut in the same ice-cold saline used for perfusion and were then transferred to a covered chamber filled with a holding solution containing (in mM): 92 NaCl, 2.5 KCl, 1.25 NaH_2_PO_4_, 30 HEPES, 25 D-glucose, 2 thiourea, 5 ascorbate, 3 pyruvate, 2 CaCl_2_, and 2 MgSO_4_. Slices were kept in this solution at room temperature for 1-2 hours before being transferred to the stage of an upright microscope for patch clamp recording. Slices were then continually perfused with an artificial CSF kept at 31°C and containing (in mM): 125 NaCl, 2.5 KCl, 1.25 NaH_2_PO_4_, 25 NaHCO_3_, 10 D-glucose, 2 CaCl_2_, and 1 MgCl_2_. All three solutions (cutting, holding, recording) were bubbled with 95% O_2_/5% CO_2_ to maintain pH at ∼7.4; all had osmolarities of 305-315 mOsm.

#### Patch Clamp Recordings

Patch clamp recordings were obtained at physiologic temperature (∼ 31 °C) under visual guidance using patch electrodes (3-7 MΩ) filled with an internal solution containing (in mM): 125 K-gluconate, 10 KCl, 4 NaCl, 10 HEPES, 4 Mg-ATP, 0.3 Tris-GTP, 7 phosphocreatine (pH 7.4, osmolarity 290 mOsm), and 0.2% neurobiotin. Before patching, fluorescence images (GFP or tdTomato) were taken with a high-resolution CCD camera to establish neuronal identity. Recordings were made in current clamp mode with bridge balance and pipette capacitance neutralization parameters set appropriately. Membrane potentials were not corrected for junction potential, which was approximately 10 mV given these solutions. Recordings with series resistances >25 MΩ, input resistances <50 MΩ, and resting potentials >-50 mV were discarded, as were recordings where any of these parameters changed by more than 20%.

To assess intrinsic excitability, neurons were held at a baseline potential of −65 mV by a DC injection, and a family of current steps (500 ms duration, −60 to 200 pA amplitude) were injected. Steps were separated by 10 s. To assess subthreshold resonance, swept-sine currents (“chirp”) between 0.5 and 12 Hz were injected (using Matlab function *chirp*), averaged, and used to extract impedance.

#### Analysis

Eighteen features were extracted from before and in the response to currents steps. (1) Membrane potential (mV), measured in the absence of a current injection. (2) Sag potential (mV), measured in response to a −60 pA step, equal to the difference between the steady-state potential and minimum potential. (3) Input resistance (MΩ), measured by the response to a −20 pA step. (4) Membrane time constant Tau (τ in ms), measured by the relaxation to a −20 pA step. (5) Rheobase (pA), the minimum current step of 500 ms duration needed to elicit an action potential. (6) Spike threshold (mV), measured from the first action potential of the rheobase current step (“first action potential”), and defined as the potential at which dV/dt crosses 20 mV/ms. (7) Spike amplitude (mV), measured from the first action potential, and defined as the difference between spike threshold and the peak of the action potential. (8) Spike width (ms), measured from the first action potential at rheobase and defined as the width at half maximum (halfway between threshold and peak). (9) Spike latency (ms), measured at rheobase current, and defined as the time difference between the start of the step and the threshold crossing of the first step. (10) Upstroke (mV/ms), the maximum value of dV/dt on the upstroke of the first action potential at rheobase (11) Downstroke (mV/ms), the minimum value of dV/dt on the downstroke of the first action potential at rheobase. (12) AHP amplitude (mV), measured following the first action potential at rheobase, defined as the difference between threshold and the minimum potential 100 ms later. (13) AHP latency (ms), the time after spike threshold is crossed by the first action potential and the AHP minimum at rheobase. (14) AHP width (ms), the time difference at half maximum of the first AHP. (15) f-I slope (Hz/pA), the slope of the initial linear section of the f-I curve. (16) Max firing rate (Hz), the maximum firing rate produced by a current step between 0 and 200 pA, across the entire 500 ms duration. (17) Adaptation index (dimensionless), for the maximal current step, the number of spikes elicited in the second half of the 500 ms step divided by the number elicited in the first 250 ms. (18) Coefficient of variation (CV) of interspike intervals, measured from the maximal current step.

### Neuronal Morphology

VP slices that contained neurobiotin filled cells were stored in 4% PFA overnight following electrophysiology recordings. Slices were then kept in PBS until immunostaining. The immunostaining protocol was identical to the immunostaining procedures described above. The primary antibody used was goat anti-ChAT (1:500, Millipore Sigma AB144P). The secondary antibodies used were donkey anti-goat A488 (Invitrogen A-11055) and streptavidin conjugated to Alexa Fluor 647 (Invitrogen S32356) to visualize neurobiotin. All images were acquired on a Nikon spinning disk confocal microscope using the following parameters: 405/488/594/647 lasers, minimal laser power (> 10%), identical exposure times for each channel, 4x averaging and 0.5 µm z-steps. All images were converted to Imaris files and the filaments tab within Imaris was used for automated neuronal tracing and analysis, including convex hull measurements and sholl analysis.

### Statistical Analysis

All statistical tests were conducted using GraphPad Prism (v. 9) or SigmaPlot (v. 14). When comparing two groups a two-tailed t-test was used and when comparing three groups a one-way ANOVA was used. Shapiro-Wilk and Smirnov-Kolmogorov tests were used to assess normality of the data. Non-parametric tests were conducted if the data failed these tests. For all statistical analysis, α was set at 0.05 and power was > 0.8.

## Supporting information

NIH Cover Letter

## Supplementary Figure Legends

**Figure S1 (supplementary to Fig 1):**
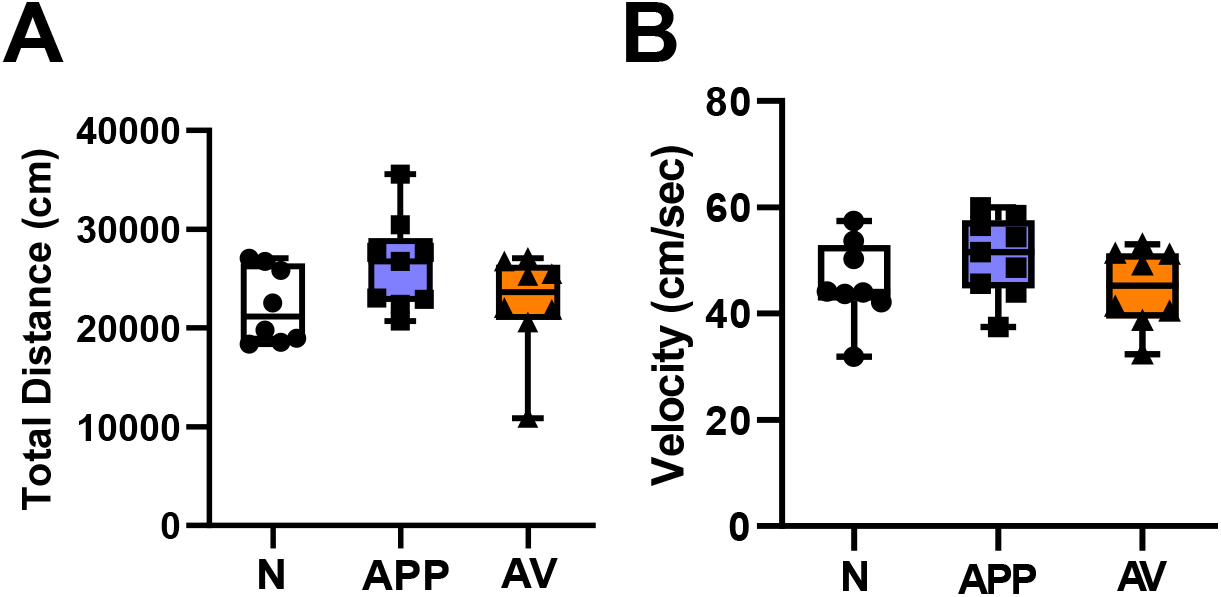
Odor induced approach and avoidance behaviors are not due to odor induced changes in locomotor activity. Neither the appetitive odor (APP) nor the aversive odor (AV) significantly altered **(A)** total distance traveled or **(B)** velocity during the odor preference test vs. a null odor stimulus (N, saline diluent).

**Figure S2 (supplementary to Fig 2):**
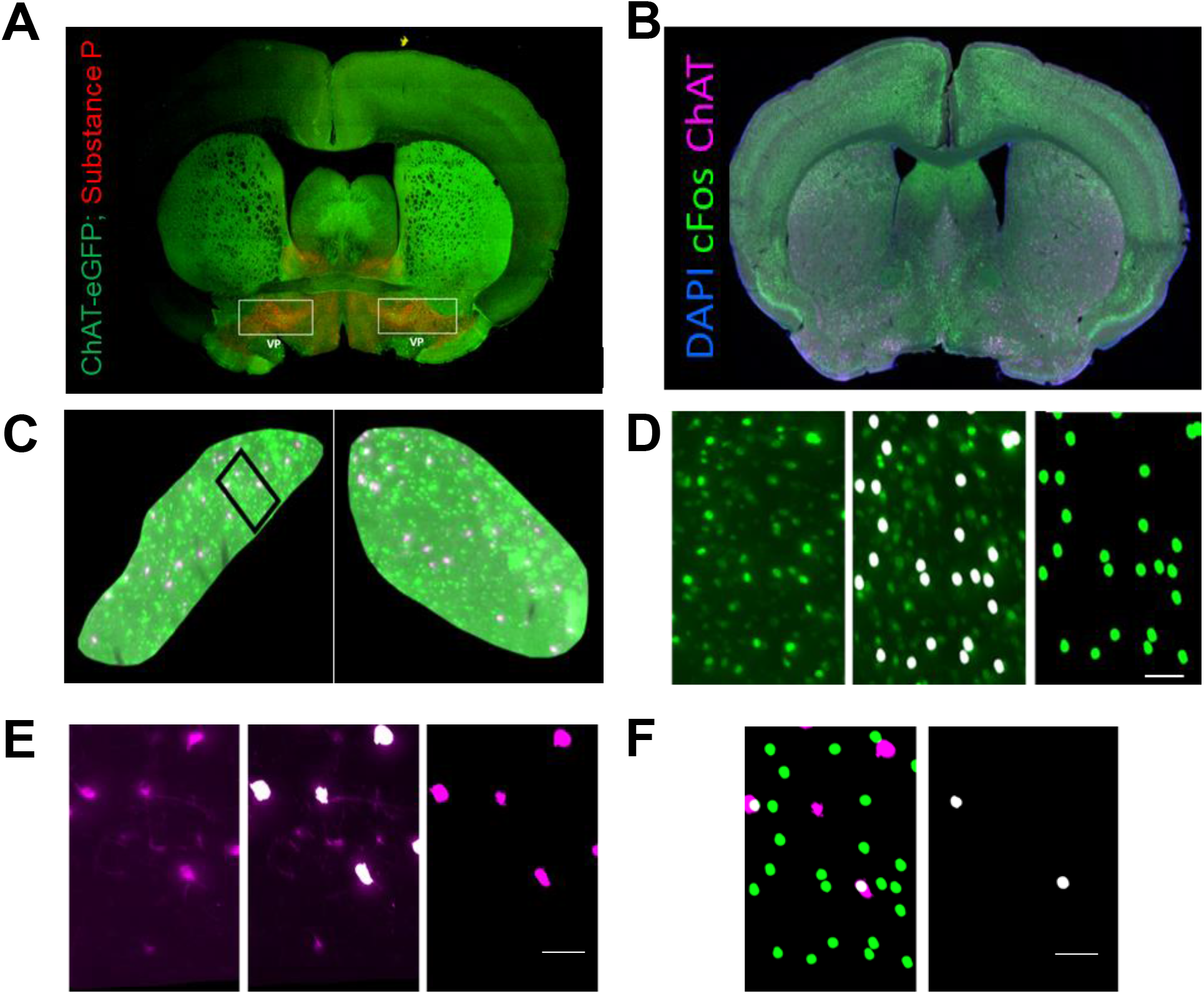
Workflow of the analysis pipeline for the detection of activated VP cholinergic neurons. **A.** Immunohistochemistry for Substance-P was conducted in a Chat-tau-eGFP mouse to assist in delineating VP borders. **B.** Following the odor preference test, tissue was collected and processed for (IHC) for ChAT (to mark cholinergic neurons) and cFos (to label activated neurons). A whole slice image was acquired using the Olympus VS200 slide scanner (20x objective, minimal exposure times for 405/488/594/647 nm channels). Files were converted to Imaris files and uploaded to Imaris software for quantification. **C.** The whole slice image is cropped and masked to contain only signal within the VP. **D.** A signal-based intensity and diameter threshold is set using the spots function in Imaris to detect cFos signal. This threshold is then used to create a new masked channel with only the cFos signal (Left: raw cFos signal, Middle: threshold for cFos signal, Right: masked cFos signal). **E.** The surfaces function in Imaris was used to set a signal-based intensity for ChAT detection. This threshold is then used to create a new masked channel containing only ChAT signal (Left: raw ChAT signal, Middle: threshold for ChAT signal, Right: masked ChAT signal) **F.** Imaris is used to detect the colocalized pixels of the two masked channels.

**Figure S3 (supplementary to Fig 4):**
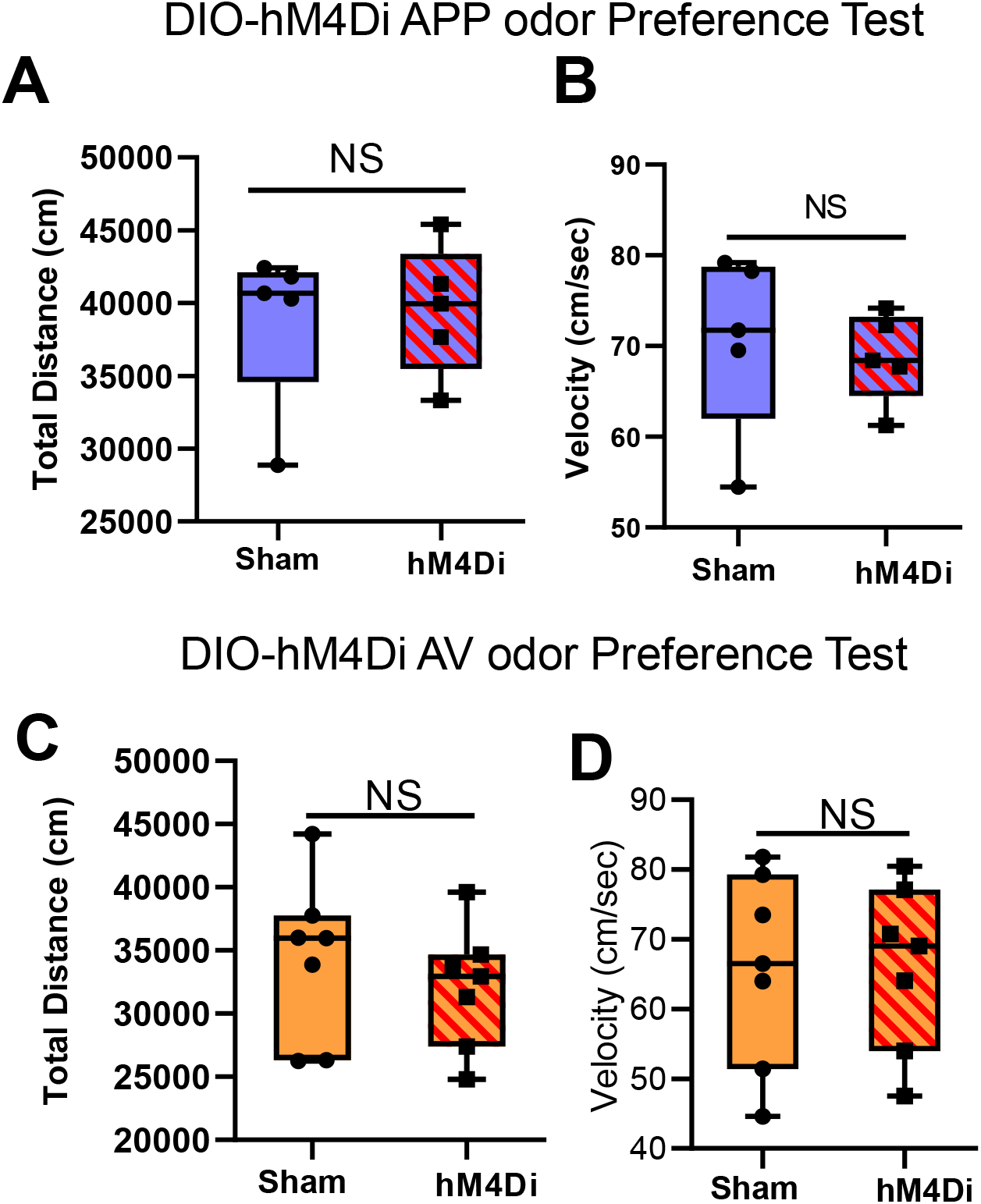
Chemogenetic inhibition of VP cholinergic neurons induce changes in odor preference but not changes in locomotor activity. To assess the effects of the inhibition of VP cholinergic neurons on approach and/or avoidance behaviors, Chat-Cre mice were injected with AAV.hSyn.DIO-hM4Di and AAV.Syn.eGFP (hM4Di experimental group) or AAV.Syn.eGFP only (Sham) in the VP. Following recovery from surgery, all mice were injected IP with 0.1 mg/kg clozapine 15-minutes prior to an odor preference test in a Y-Maze. In the odor preference test, mice were allowed access to two arms of the Y-Maze (appetitive (APP) odor vs. saline **(A & B)** or aversive (AV) odor vs. saline **(C & D)**). Time spent in each arm, as well as locomotor activity (distance traveled and velocity) were assessed. Top = DIO-hM4Di APP odor preference test. There is no difference between mice in the sham group and mice that express DIO-hM4Din **(A)** distance traveled or **(B)** velocity during the APP odor preference test. Bottom = DIO-hM4Di AV odor preference test. There is no difference between mice in the sham group and mice that express DIO-hM4Di in **(A)** distance traveled or **(B)** velocity during the AV odor preference test.

**Figure S4 (supplementary to Fig 5 & 6):**
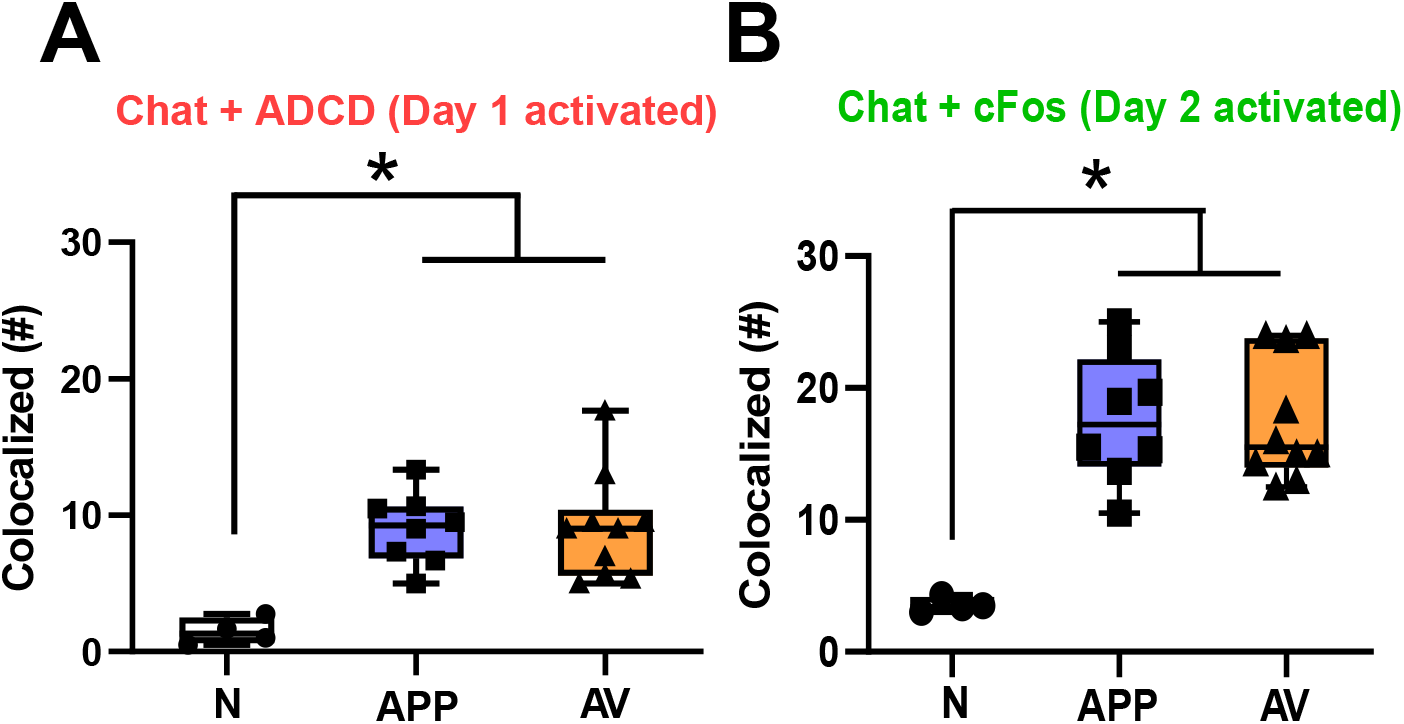
The order of odor presentation does not affect the total number of VP cholinergic neurons activated. Odor exposure (either appetitive odor (APP) or aversive odor (AV)) significantly increases the number of activated VP cholinergic neurons compared to a null odor stimulus (N, saline diluent). Regardless of order of odor presentation (i.e., (**A**) Day 1 vs. (**B**) Day 2) and method in which activated VP cholinergic neurons are assayed (i.e., (**A**) ADCD vs. (**B**) ChAT and cFos IHC), both odors (APP or AV) significantly increase the number of activated VP Cholinergic neurons vs. a null odor stimulus. * *p* < 0.05.

**Figure S5 (supplementary to Fig 5 & 6):**
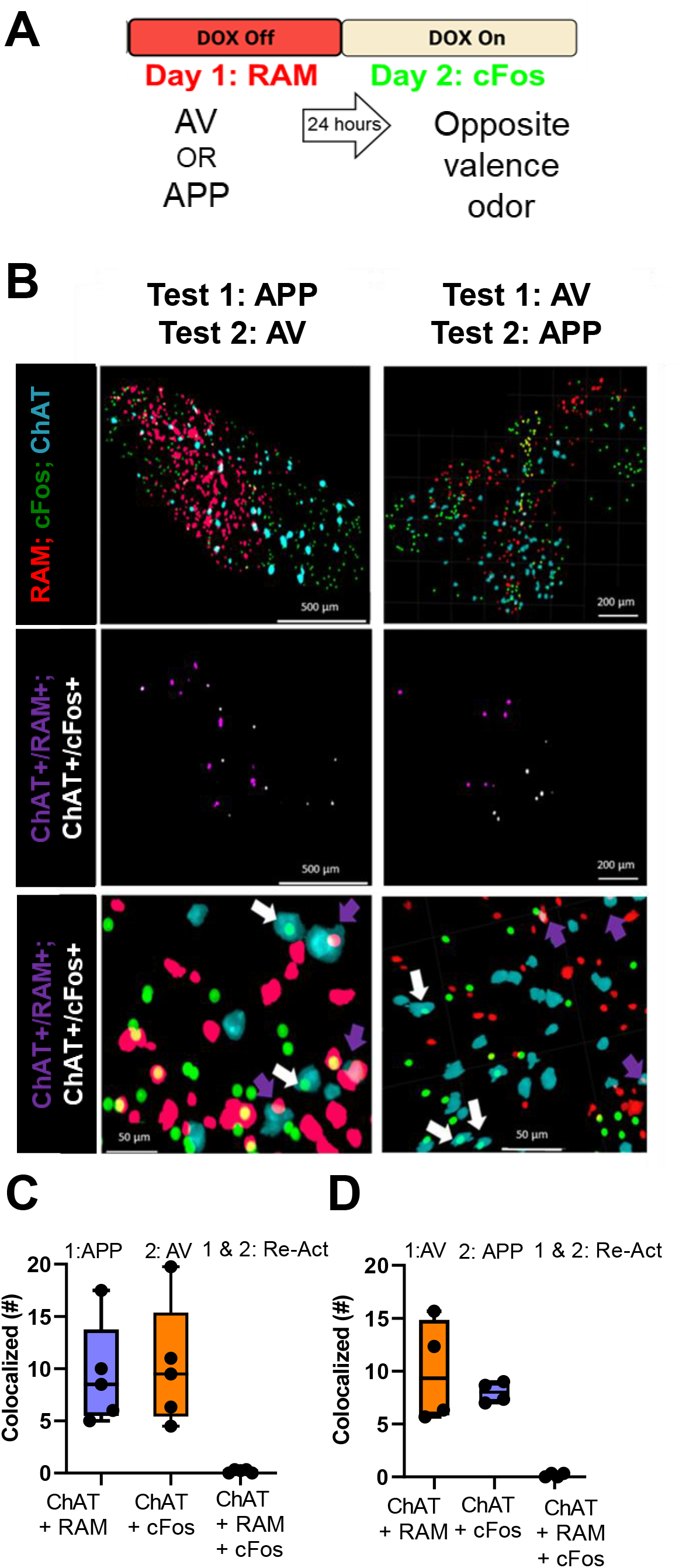
Experiments using the robust activity marker (RAM) confirm ADCD and cFos labeling experiments. **A.** A distinct activity-dependent viral vector was used to verify results from ADCD and cFos labeling experiments. The robust activity marker (RAM) viral vector utilizes a synthetic activity-dependent promoter and a Tet-Off system to label activated neurons. RAM was injected in the VP of wild-type C57/BL6J mice. The RAM construct was used in conjunction with ChAT and cFos IHC to label activated VP cholinergic neurons in two distinct contexts. The behavioral paradigm used was identical to the protocol used for ADCD and cFos labeling experiments. Representative images from RAM experiments showing RAM positive (activated neurons on Day 1), cFos positive (activated cells on Day 2), ChAT (cholinergic marker), and the colocalization of RAM + ChAT and cFos + ChAT. **B. & C.** Confirming results from the ADCD and cFos labeling experiments (in Fig 6), mice exposed to a different odor on Day 2 exhibited no overlap between ChAT+/RAM+ and ChAT+/cFos+ neurons.

**Figure S6 (supplementary to Fig 5 & 6):**
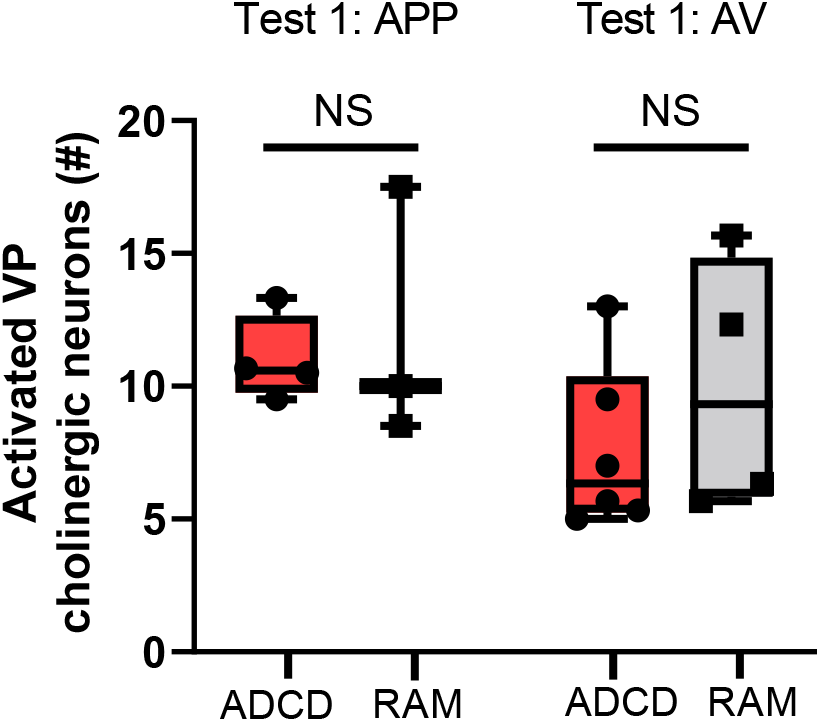
ADCD and RAM are comparable in labeling activated VP cholinergic neurons. For ADCD experiments, Chat-Cre x Fos-tTA/GFP mice were injected with ADCD in the VP. For Test 1, ADCD was used to assess the number of activated VP cholinergic neurons. On Test 2, the colocalization of ChAT and cFos-GFP (examined using IHC) was used to assess the number of activated cholinergic neurons (see Fig 5 and Fig 6 for details). For RAM experiments, WT mice were injected with RAM in the VP. For Test 1, the number of RAM+ neurons co-labeled with ChAT IHC was used to assess the number of activated VP cholinergic neurons. For Test 2, the colocalization of ChAT and cFos (examined using IHC) was used to assess the number of activated VP cholinergic neurons (see Fig S5 for details). Regardless of order of odor presentation (Left = appetitive odor (APP), Right = aversive odor (AV)), ADCD and RAM labeled similar number of activated VP cholinergic neurons.

**Figure S7 (supplementary to Fig 7):**
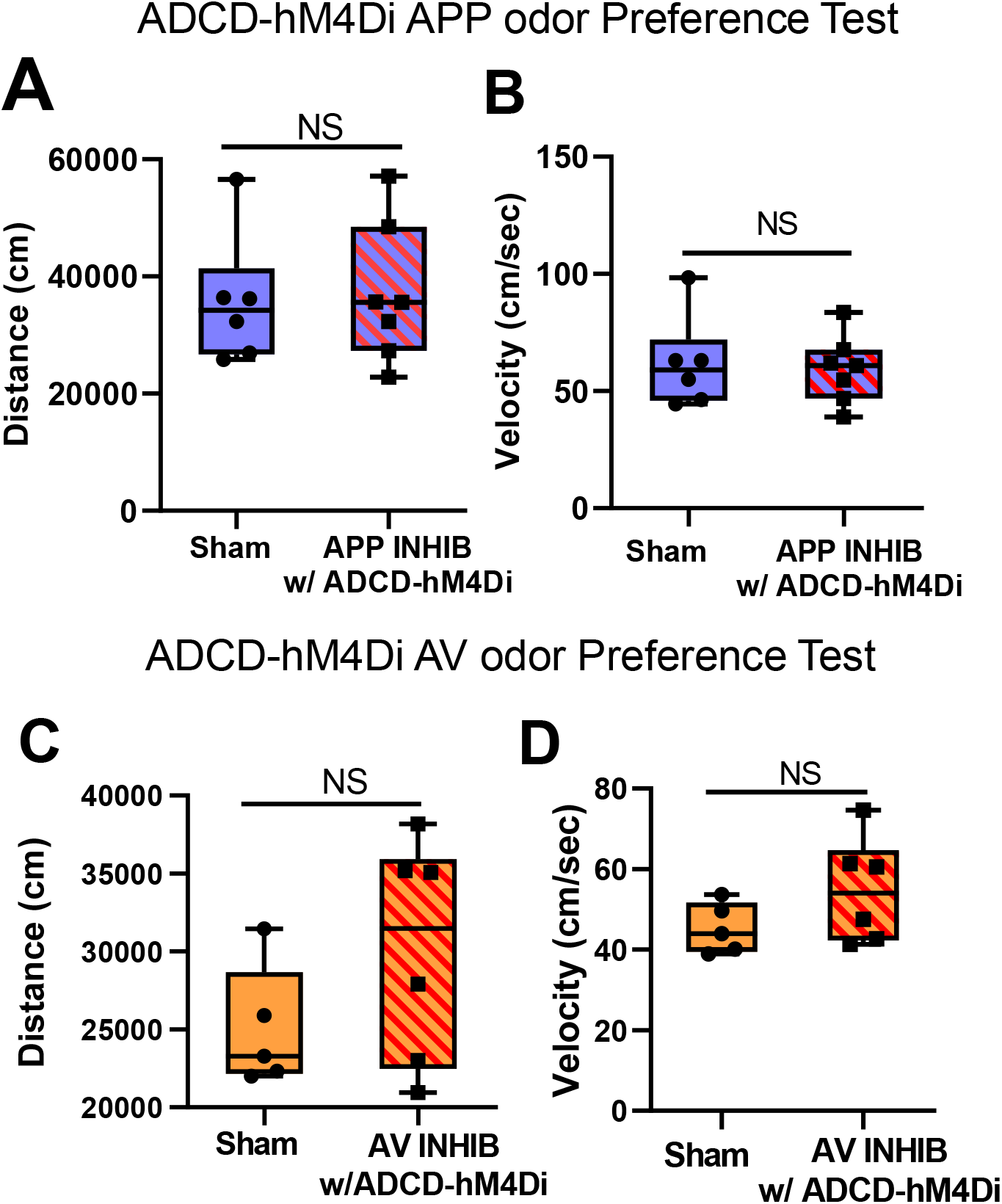
ADCD-hM4di induced changes in odor preference are not due to changes in locomotor activity. To assess the effects of the inhibition of previously activated VP cholinergic neurons on approach and/or avoidance behaviors, Chat-Cre x Fos-tTA/GFP mice were injected with ADCD and AAV.Syn.eGFP (experimental group), or AAV.Syn.eGFP only (Sham) in the VP (see Fig 10 legend and methods for details). Following recovery from surgery, mice were habituated in the Y-Maze (2 x 10 min) and taken off a DOX diet. Approximately 24-hours later, mice were exposed to an odor (either appetitive (APP) or aversive (AV)) in one arm of the Y-Maze. Following odor exposure, mice were placed on a DOX diet to prevent further expression of ADCD. Approximately 24-hours later, all mice were injected IP with 0.1 mg/kg clozapine 15-minutes prior to an odor preference test in a Y-Maze. In the odor preference test, mice were allowed access to two arms of the Y-Maze (previously exposed odor, either APP or AV, vs. saline). Time spent in each arm, as well as locomotor activity (distance traveled and velocity) were assessed. Top = ADCD-hM4Di appetitive (APP) odor preference test. There is no difference between mice in the sham group and mice that express ADCD-hM4Di in **(A)** distance traveled or **(B)** velocity during the APP odor preference test. Bottom = ADCD-hM4Di aversive (AV) odor preference test. There is no difference between mice in the sham group and mice that express ADCD-hM4Di in **(C)** distance traveled or **(D)** velocity during the AV odor preference test.

**Figure S8 (supplementary to Fig 9):**
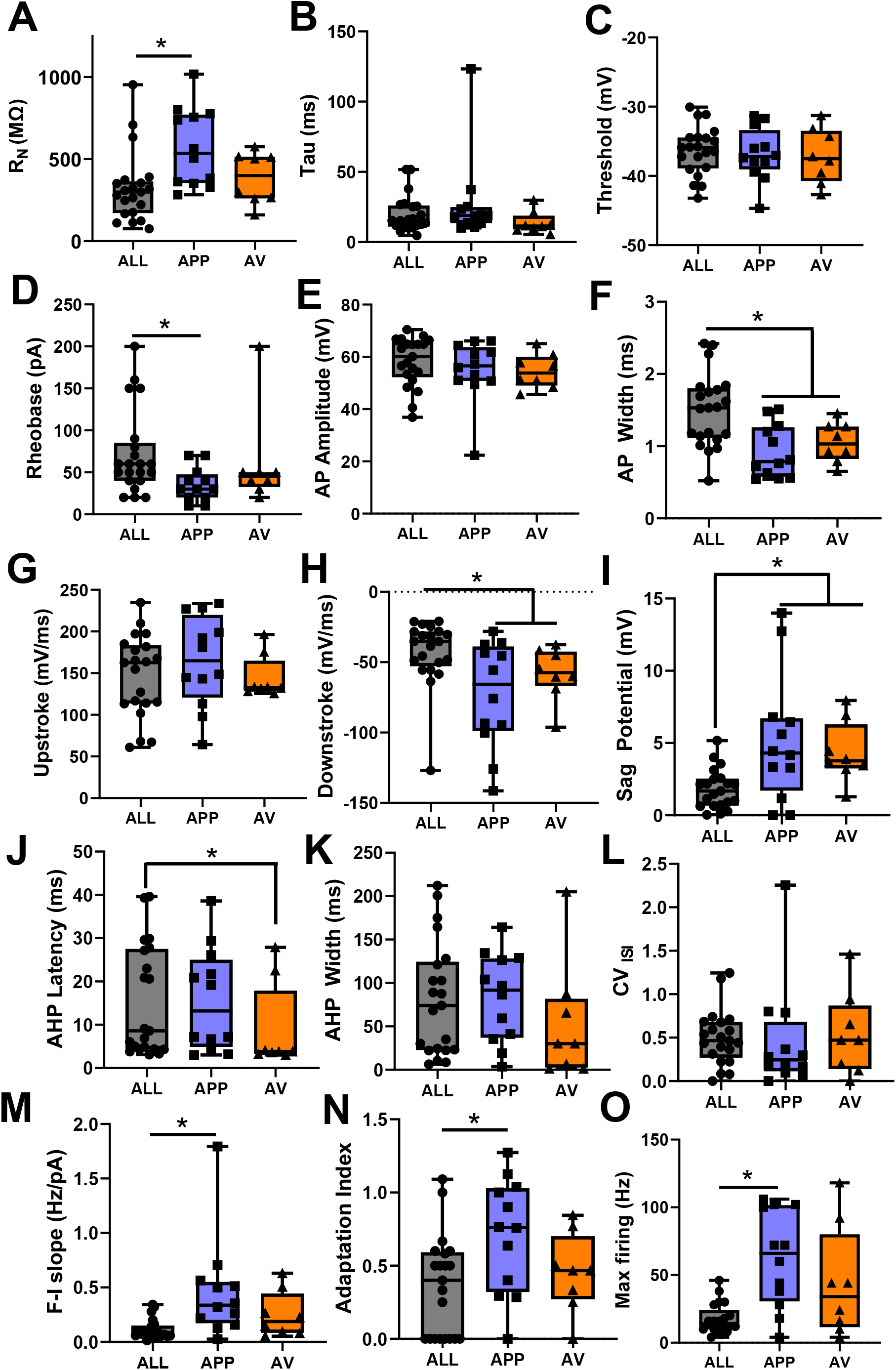
Comparison of the electrophysiological properties of appetitive (APP) vs. AV (AV) VP cholinergic neurons demonstrates that they are largely similar to one another and to the overall population of VP cholinergic neurons (see Fig 8 for differences). The majority of both passive and active membrane properties are the same whether the recordings are from APP or AV activated VP cholinergic neurons. Similarities between the two are observed in **(A)** input resistance, **(B)** tau, **(C)** threshold, **(D)** rheobase, **(E)** action potential amplitude, **(F)** action potential width, **(G)** action potential upstroke, **(H)** actional potential downstroke, **(I)** sag potential, **(J)** afterhyperpolarization latency, **(K)** afterhyperpolarization width, **(L)** coefficient variation**, (M)** frequency-current slope, **(N)** adaptation index and **(O)** max firing. Both APP and AV activated VP cholinergic neurons significantly differ from ALL VP cholinergic neurons in **(F)** action potential width and **(I)** sag potential. APP odor activated VP cholinergic neurons significantly differ from ALL VP cholinergic neurons in **(A)** input resistance, **(D)** rheobase, **(M)** frequency-current slope, **(N)** adaptation index and **(O)** max firing. AV odor activated VP cholinergic neurons significantly differ from ALL VP cholinergic neurons in **(J)** afterhyperpolarization latency.

**Figure S9 (supplementary to Figure 10):**
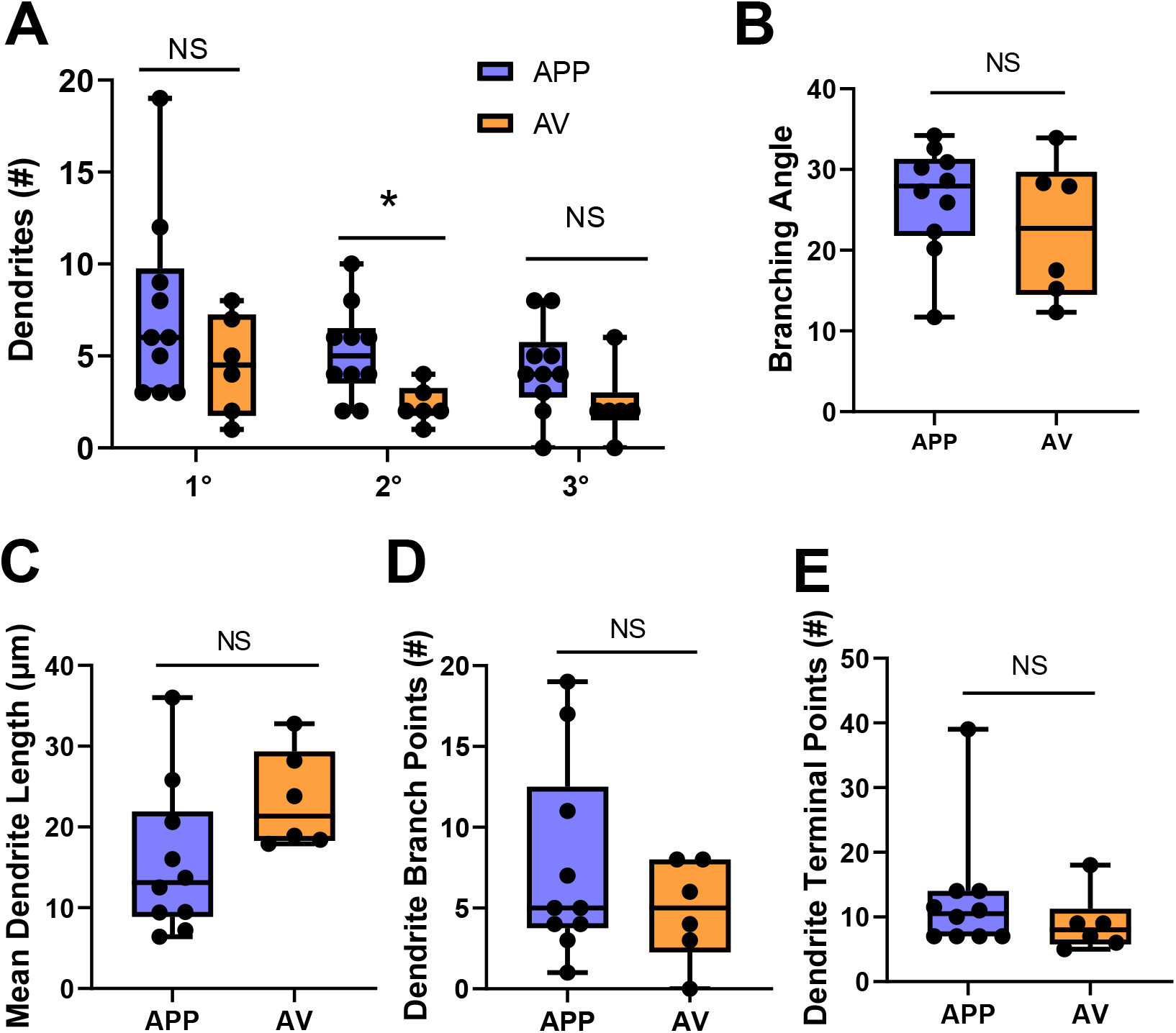
Additional morphological properties between appetitive odor activated (APP) and aversive odor activated (AV) VP cholinergic neurons. **A.** Assay of the number of 1°, 2°, and 3° dendrites revealed statistically significant differences between APP vs. AV VP cholinergic neurons for only 2^nd^ order dendrites. There was no significant difference between groups in the number of primary or tertiary dendrites. The number of secondary dendrites was significantly higher in appetitive odor (APP) vs. aversive odor (AV) activated VP cholinergic neurons (* *p* < 0.05). Despite differences in proximal complexity (see Fig 9), there was no significant differences between groups in **(B)** branching angle, **(C)** mean dendrite length, **(D)** number of dendrite branch points or **(E)** number of dendrite terminal points.

